# Lack of Oncomodulin Increases ATP-Dependent Calcium Signaling and Susceptibility to Noise in Adult Mice

**DOI:** 10.1101/2024.06.10.598303

**Authors:** Yang Yang, Jing-Yi Jeng, Kaitlin Murtha, Leslie K. Climer, Federico Ceriani, Weintari D. Sese, Aubrey J. Hornak, Kiah C. Sleiman, Ian Stahl, M. Charles Liberman, Walter Marcotti, Dwayne D. Simmons

## Abstract

Tight regulation of Ca^2+^ is crucial for the function of cochlear outer hair cells (OHCs). Dysregulation of Ca^2+^ homeostasis in OHCs is associated with impaired hearing and contributes to increased vulnerability to insults such as noise exposure. Ca^2+^ signaling in developing OHCs is modulated by oncomodulin (OCM), an EF-hand calcium-binding protein. Here, we investigated whether the lack of OCM disrupts the control of intracellular Ca^2+^ in mature OHCs, and influences vulnerability to acoustic injury. Using young adult CBA/CaJ mice, we found that OHCs from *Ocm*-knockout (*Ocm^-/-^*) mice showed normal biophysical profiles, electromotile responses, and synaptic innervation compared to littermate controls. Moderate noise exposure (95 dB SPL, 2 hrs) caused temporary threshold shifts in *Ocm*^+/+^ and *Ocm*^-/-^ mice. However, while *Ocm*^+/+^ fully recovered thresholds 2 weeks after noise exposure, *Ocm*^-/-^ mice showed permanent threshold shifts. Additionally, *Ocm*^-/-^ mice had auditory brainstem responses with highly variable latencies and amplitudes both before and after noise exposure compared to *Ocm*^+/+^ mice. Using a genetically encoded Ca^2+^ sensor (GCaMP6s) expressed in *Ocm*^+/+^ and *Ocm*^-/-^ OHCs, we found that prolonged noise exposure (95 dB SPL, 9 hrs) significantly increased GCaMP6s fluorescence, ATP-induced Ca^2+^ signaling and also caused greater threshold shifts in *Ocm*^-/-^ compared to *Ocm*^+/+^ OHCs. However, prolonged noise exposure had no significant change in the number of presynaptic OHC ribbons in either *Ocm*^+/+^ or *Ocm*^-/-^ mice. We assessed whether the ATP-induced responses were due to changes in P2X2 receptor expression. Prior to noise exposure, P2X2 expression was higher in the cochlea of *Ocm*^-/-^ mice compared to *Ocm*^+/+^ mice. Following prolonged noise, P2X2 receptors were upregulated in the cochlea of *Ocm*^+/+^ but not in the *Ocm*^-/-^ mice, which retained their pre-noise expression level. We propose that the lack of OCM increases susceptibility to cochlear pathology and that purinergic signaling and dysregulation of cytosolic Ca^2+^ homeostasis likely contribute to early onset hearing loss in the *Ocm*^-/-^ mice.

## Introduction

The tight regulation of Ca^2+^ is crucial for cochlear sensory cells. In response to sound stimulation, Ca^2+^ influx and Ca^2+^-induced Ca^2+^ release (CICR) from the endoplasmic reticulum significantly increase intracellular Ca^2+^ in cochlear sensory cells. Factors that cause hearing impairment, such as ototoxic drugs, acoustic overstimulation, and aging, are associated with dysregulation of Ca^2+^ homeostasis (Fridberger & Ulfendahl, 1996; Liu & Fechter, 1997; Kidd Iii & Bao, 2012; Richard *et al*., 2023). Two types of specialized sensory cells in the mammalian cochlea, inner hair cells (IHCs) and outer hair cells (OHCs), are involved in the transduction of sound into electrical responses (Dallos, 1992). The OHCs are largely responsible for the sensitivity and tuning, whereas the IHCs transmit auditory signals to the brain via the auditory nerve. One of the hallmarks of cochlear injury following noise exposure is loss of OHCs. However, comparatively little is known about the endogenous mechanisms necessary to protect OHCs from the damaging effects of noise exposure. Shortly after acoustic overstimulation, intracellular Ca^2+^ concentration is increased in auditory hair cells, especially OHCs (Fridberger *et al*., 1998a; Zuo *et al*., 2008), which leads to their loss (Fridberger *et al*., 1998a; Orrenius *et al*., 2003; Esterberg *et al*., 2013). Thus, effective regulation of Ca^2+^ is essential for the responses of OHCs to noise.

Although evidence strongly suggests that Ca^2+^ overload can trigger apoptotic or necrotic cell death, and thus contribute to noise-induced damage, molecular models for Ca^2+^ overload are lacking. In OHCs, Ca^2+^ homeostasis depends upon the balance of Ca^2+^ influx, extrusion, and buffering. Oncomodulin (OCM) is a small, 12-kDa, EF-hand calcium-binding protein (CaBP) that is predominantly expressed in OHCs and functions differently from other EF-hand CaBPs (Pangrsic *et al*., 2015; Climer *et al*., 2019; Murtha *et al*., 2022). OCM is the β isoform of parvalbumin and shares at least 53% sequence identity with α-parvalbumin (PVALB). In adult mice, targeted deletion of OCM (*Ocm*^-/-^) is associated with early progressive hearing loss and to date, is the only EF-hand CaBP in OHCs associated with hearing loss. *Ocm*^-/-^ mice have earlier onset of age-related threshold shifts, independent of genetic background, which indicates the importance of OCM in the long-term maintenance of OHC function (Tong *et al*., 2016; Climer *et al*., 2021). Recent studies have shown that the absence of OCM increases Ca^2+^ signaling in developing OHCs, and reveal that OCM plays a significant role in the synchronization of Ca^2+^ activity during cochlear maturation (Murtha *et al*., 2022; Yang *et al*., 2023). In Murtha, Yang, et al. (2022), we found that the onset of OCM expression alters the level of Ca^2+^ signals in OHCs before hearing onset. While the expression of other CaBPs changed in response to the loss of OCM, these changes did not compensate for the loss of OCM function. These results support the idea that OCM plays a unique and indispensable role as a Ca^2+^ buffer in developing OHCs. Our previous findings also revealed higher levels of spontaneous Ca^2+^ activity, upregulation of purinergic receptors in OHCs, and higher synchronization of Ca^2+^ activity in pre-hearing *Ocm*^-/-^ mice compared to wild-type mice (Yang et al. 2023). Additionally, the maturation of OHC ribbons in *Ocm*^-/-^ mice was delayed, leading to an increased number of ribbons and afferent fibers on OHCs before hearing onset. Given that OCM can modulate Ca^2+^ signaling and synchronization in OHCs during the pre-hearing period, and is essential for maintaining hearing in young adults, we hypothesize that the lack of OCM in adult mice increases OHC vulnerability to noise.

In a recent study (Murtha et al. 2024), we show that the absence of *Ocm* leads to an increased susceptibility to OHC loss after exposure to high-level, broadband noise. Moreover, in *Ocm^-/-^* OHCs, we observed changes in mitochondrial protein expression, both with and without noise exposure. In the present study, we used young adult *Ocm*^-/-^ and age-matched controls to study the biophysical properties, morphological characteristics, and purinergic receptor expression in OHCs, and their susceptibility to permanent hearing threshold shifts. Although the OHCs of young adult *Ocm*^-/-^ mice exhibited normal biophysical properties and electromotile responses, and afferent innervation, the latency and amplitude of wave I auditory brainstem response showed greater variability. In response to moderate noise exposures, *Ocm*^-/-^ mice showed larger threshold shifts and more variable amplitude and latency compared to litermate controls. Without noise stimulation, adult OHCs in *Ocm*^-/-^ mice also had higher purinergic P2X2 receptor expression and larger Ca^2+^ response induced by extracellular ATP. However, in response to noise exposure, P2X2 expression increased in the OHCs from *Ocm*^+/+^, but not in *Ocm*^-/-^ cochlea. We conclude that the increased vulnerability to noise in *Ocm*^-/-^ mice may be associated with pre-existing, elevated levels of both Ca^2+^ signaling and purinergic receptor expression, and ultimately leads to an early onset of age-related hearing loss observed in our previous reports (Tong *et al*., 2016; Climer *et al*., 2021).

## Materials and Methods

### Ethical approval

All procedures performed in the USA were approved by Baylor University and Mass Eye and Ear Institutional Animal Care and Use Committees (IACUC) as established by the US Public Health Service and performed in compliance with the National Institutes of Health animal care guidelines. Procedures performed in the UK and licensed by the Home Office under the Animals (Scientific Procedures) Act 1986 were approved by the University of Sheffield Ethical Review Committee.

### Animals

Animals were bred in animal facilities at Baylor University, Mass Eye and Ear, and the University of Sheffield. The original *Ocm* mutant mouse (C57Bl/6 ActbCre; Ocmflox/flox) (Tong *et al*., 2016) was backcrossed onto the CBA/CaJ and the CBA/CaH background to minimize or eliminate any confounding effects of Cdh23ahl mutation, which is linked to hearing loss in the adult (Liu *et al*., 2007; Climer *et al*., 2021; Murtha *et al*., 2022). Studies using CBA/CaH mice were performed at the University of Sheffield, where mice were sacrificed via cervical dislocation in accordance with UK Home Office regulations.

We also generated mice with a genetically encoded, tissue-specific Ca^2+^ sensor (*Atoh1*-GCaMP6s) in *Ocm* wild-type (GCaMP6s-*Ocm*^+/+^) and *Ocm* knockout (GCaMPs-*Ocm*^-/-^) mice (Yang *et al*., 2023). We first crossed *Ocm* wild-type (*Ocm^+/+^*) and *Ocm* knockout (*Ocm^-/-^*) mice with B6;129S-*Gt(ROSA)26Sor^tm96.1(CAG-GCaMP6s)Hze^*/J, Ai96(RCL-GCaMP6s) or Ai96, which contains a floxed-STOP cassette preventing transcription of the GCaMP6 slow variant Ca^2+^ indicator. These mice were then crossed with knock-in transgenic mice expressing Cre recombined from the *Atoh1* locus (Chen *et al*., 2013). *Atoh1*-driven Cre GCaMP6s mice showed tissue-specific expression of endogenous green fluorescence (Yang *et al*., 2010; Cox *et al*., 2012; Mulvaney & Dabdoub, 2012).

### Single-cell electrophysiology

For electrophysiological recordings, OHCs from CBA/CaH control (*Ocm^+/-^*) and *Ocm*-knockout (*Ocm^-/-^*) mice of both sexes were acutely dissected at 1 - 2 months. Cochleae were isolated from the inner ear as previously reported (Marcotti & Kros, 1999; Ceriani *et al*., 2019; Jeng *et al*., 2020; Jeng *et al*., 2021; Murtha *et al*., 2022) using an extracellular solution composed of (in mM): 131 NaCl, 5.8 KCl, 1.3 CaCl_2_, 0.9 MgCl_2_, 0.7 NaH_2_PO_4_, 5.6 D-glucose, 10 HEPES-NaOH. Sodium pyruvate (2 mM), amino acids, and vitamins (Thermo Fisher Scientific, UK). The pH was adjusted to 7.5 (osmolality ∼308 mmol kg^-1^). The dissected cochleae were fixed at the bottom of the recording chamber by a nylon-meshed silver ring and perfused with the above extracellular solution. OHCs were viewed using an upright microscope (Olympus BX51) equipped with Nomarski Differential Interface Contrast (DIC) optics with a 60x water immersion objective and 15x eyepieces.

Recordings were performed at room temperature (21-24°C) using an Optopatch amplifier (Cairn Research Ltd, UK). Patch pipettes were pulled from soda glass capillaries, and the shank of the electrode was coated with surf wax (2–3 MΩ). Current and voltage responses were measured using the following intracellular solution (in mM): 145 KCl, 3 MgCl_2_, 1 EGTA-KOH, 5 Na_2_ATP, 5 HEPES-KOH, 10 sodium phosphocreatine (pH adjusted to 7.28 with KOH; osmolality was 294 mmol kg^−1^). Voltage clamp protocols were performed at a holding potential of −84 mV. Data acquisition was performed using pClamp software (Axon Instruments, Union City, CA, USA) using Digidata. Data were filtered at 2.5 kHz (8-pole Bessel), sampled at 5 kHz, and stored on the computer. Offline data analysis was performed using Origin software (OriginLab, Northampton, MA, USA). Membrane potentials reported were corrected for the uncompensated residual series resistance (*R*_s_) and the liquid junction potential (LJP), which was -4 mV, measured between electrode and bath solutions.

### Electromotile response

Electromotility was estimated in OHCs at room temperature (∼22°C) by applying a depolarizing voltage step from the holding potential of -64 mV to +56 mV and recorded using a CCD camera (Thorlabs DCU224M). The camera was attached to a microscope (Olympus), equipped with a X60 water immersion objective (Olympus LUMPlanFL N). The acquired images were stack-sliced along a vertical axis of each OHC, and the contraction was measured on the image stack as length change of the cell. All images were analyzed in ImageJ and the measurements were calibrated using a stage graticule.

### Cochlear Function Assays

For measurement of auditory brainstem responses (ABRs) and distortion product otoacoustic emissions (DPOAEs), adult mice were anesthetized with xylazine (20 mg/kg, i.p.) and ketamine (100 mg/kg, i.p.). Acoustic stimuli were delivered using a custom acoustic assembly (Maison *et al*., 2012). Briefly, two electrostatic earphones (CUI CDMG15008-03A) were used to generate primary tones and a Knowles miniature microphone (EK-3103) was used to record ear-canal sound pressure. Stimuli were generated digitally with 4 μs sampling. Ear-canal sound pressure and electrode voltage were amplified and digitally sampled at 20 μs for analysis of response amplitudes. The acoustic stimuli and physiological responses were digitized by a National Instruments PXI system with 24 - bit sound cards running custom LabVIEW software. ABRs were recorded via 30-gauge platinum electrodes, inserted subdermally, adjacent to the pinna incision and at the vertex of the skull, with a ground near the tail. Tone-pip stimuli were presented at half-octave frequency intervals from 5.6 - 45.2 kHz. At each frequency, a level series was presented, in either 5 or 10 dB steps, from below threshold to 80 dB SPL, with up to 1024 stimuli averaged per step. Sound-evoked responses were amplified 10,000x through a 0.3 - 3 kHz bandpass filter. ABR threshold was defined as the lowest stimulus level to generate a waveform that increased in amplitude and decreased in latency as the stimulus level increased. The amplitude and latencies of the ABR waveform potentials were separately measured for the wave I (P1 - N1) (Ceriani, 2024). For measurement of DPOAEs at 2f1 - f2, the primary tones were set so that the frequency ratio (f2/f1) was 1.2 and so that f2 level was 10 dB below f1 level. For each f2/f1 primary pair, levels were swept in 5 or 10 dB steps from 10 dB SPL to 80 dB SPL (for f2). At each level, both waveform and spectral averaging were used to increase the signal-to-noise ratio of the recorded ear-canal sound pressure, and the amplitude of the DPOAE at 2f1 - f2 was extracted from the averaged spectra, along with the noise floor at nearby points in the spectrum. Iso-response curves were interpolated from plots of DPOAE amplitude vs. sound level. The threshold was defined as the f2 level required to produce a DPOAE at 0 dB SPL. Right ears were used for all hearing tests. For the total threshold shift of ABR and DPOAE, the total threshold shifts across all frequencies for each animal were calculated and then averaged across all animals.

### Noise exposure

Two different experimental approaches were used to assess the effects of noise exposure. The first approach assessed temporary and permanent threshold shifts, while the second approach assessed the immediate effects of longer noise exposure. 4 - 6 wks old *Ocm*^+/+^ and *Ocm*^-/-^ CBA/CaJ mice and 3 - 4 wks old GCaMP6s - *Ocm*^+/+^ and - *Ocm*^-/-^ were randomly assigned to either noise-exposed or control groups. Hearing thresholds of age-matched mice were measured 24 hrs prior to noise exposure. Unexposed mice served as controls. CBA/CaJ mice were exposed to an octave band noise for 2 hrs then cochlear function was assessed at 6 hrs and 2 wks later, followed by cochlear fixation and tissue harvest. For GCaMP6s mice, exposure was to broadband noise at 95 dB SPL for 9 hrs. Cochlear function was assessed 1 hr following noise exposure followed by tissue harvesting.

For both groups, awake and unrestrained mice were placed in individual sections of a wire-mesh cage and exposed to either octave band noise (8-16 kHz) for 2 hrs at 95.0 dB SPL, or broadband noise for 9 hrs at 95 dB SPL. The booth is equipped with a JBL2446H compression driver coupled to an exponential horn. Noise levels were calibrated and monitored either with a 1⁄4′′ Brüel and Kjær condenser microphone or a sound-level meter. The noise level varied by less than 2 dB over the duration of the exposures. Food and water were provided in a petri dish during 9 hrs of noise exposure.

### Tissue preparation

Cochleae were harvested from 3 - 4 wks old *Ocm*^+/+^ or *Ocm*^-/-^ mice of either sex. After decapitation, apical coil OHCs were dissected from the organ of Corti in an extracellular medium composed of (in mM): 136.8 NaCl, 5.4 KCl, 0.4 KH2PO4, 0.3 Na2HPO4, 0.8 MgSO4, 1.3 CaCl2, 4.2 NaHCO3, 5 HEPES and 5.6 glucose. The pH was adjusted to 7.4 - 7.5 (osmolality ∼306 mmol/kg). The apical coil was then transferred to a small microscope chamber with nylon mesh fixed to a stainless-steel ring on the bottom and visualized using an upright microscope (Leica, DM 6000 FS, Germany) with a water immersion objective (Leica, Germany).

### Confocal Ca^2+^ imaging

Ca^2+^ signals from GCaMP6s were recorded at room temperature with 480nm excitation wavelength (X-Light V2 spinning disk confocal, 89 North Inc, PRIME95B Photometrics Cooled sCMOS with 95% QE, Teledyne Photometric). Images were taken using VisiView (VISITRON, USA) and analyzed offline using ImageJ (NIH). Ca^2+^ signals were measured as relative changes in fluorescence emission intensity (ΔF/F_0_) and calculated by MATLAB. ΔF = F - F_0_, where F is fluorescence at time t and F_0_ is the fluorescence at the onset of the recording.

For inducing Ca^2+^ transients, 100 µl of ATP depolarization solution was perfused into a microscope chamber containing 300 µl extracellular solution (1:4 dilution, 100 µM ATP final concentration in the chamber) at 8 s after the imaging started, followed by a 2 min perfusion with the extracellular solution to help with equilibration and wash out. Only the fluorescence signals from OHCs that reached maximum intensity between 8-60 s were calculated. For the P2X antagonist experiments, the blocker, PPADS (100 μM), was added to the explant right before ATP application. Each GCaMP6s fluorescence recording includes 1500 frames taken at 15 frames per second from 873 ×873 pixels region. After background subtraction, the time course of Ca^2+^ changes in activated OHCs was computed as pixel averages of a circle region of interest (ROI) using ImageJ.

### Genotyping and qRT-PCR

DNA was extracted from mice tail samples using Extract-N-Amp™ Tissue PCR Kit (Sigma, USA). PCR primers used for genotyping are listed below: *Atoh1*-Cre primer pair forward: 5’-CCGGCAGAGTTTACAGAAGC-3’, reverse: 5’-ATG TTT AGC TGG CCC AAA TG-3’; Cre control primer pair forward: 5’-CTA GGC CAC AGA ATT GAA AGA TCT-3’; reverse: 5’-GTA GGT GGA AAT TCT AGC ATC ATC C-3’; GCaMP6s primer pair forward: 5’-ACG AGT CGG ATC TCC CTT TG - 3’; reverse: 5’-AGA CTG CCT TGG GAA AAG CG - 3’; *Ocm* primer pair forward: 5’-CTC CAC ACT TCA CCA AGC AG - 3’, reverse: 5’-TTT CAT GTT CAG GGA TCA AGT G - 3’; *Ocm* deletion primer pair forward: 5’-CTC CAC ACT TCA CCA AGC AG - 3’, reverse: 5’-GCT TGG GGA CCC CCT GTC TTC A - 3’.

Cochleae were acutely dissected after anesthesia and transferred to the lysis buffer. Total RNA was extracted using RNeasy plus Micro kits (QIAGEN, USA). iScript™ Advanced cDNA Synthesis Kit (BIO-RAD, USA) was used for reverse transcription.

qRT-PCR was performed using the SYBR Green PCR Master Mix Kit (BIO-RAD, USA) as previously described (Murtha *et al*., 2022). Briefly, the *b2m* gene was used as a reference gene (Melgar-Rojas *et al*., 2015). Quantification of expression (fold change) from the Cq data was calculated following the ΔΔCq method (Schmittgen & Livak, 2008), and normalized to the Cq value of *Ocm*^+/+^. 2^−ΔΔCq^ was calculated to represent the relative expression (fold change). Primers used for qRT-PCR are as follows: *b2m* forward 5’-TGGTCTTTCTGGTGCTTGTC-3’ and reverse 5’-GGG TGG AAC TGT GTT ACG TAG-3’; *P2RX2* forward 5’-GCG TTC TGG GAC TAC GAG AC -3’ and reverse 5’-ACG TAC CAC ACG AAG TAA AGC -3’; *Ocm* primer pair forward: 5’-CTC CAC ACT TCA CCA AGC AG - 3’, reverse: 5’-TTT CAT GTT CAG GGA TCA AGT G - 3’ (PrimerBank ID 27544798a1).

### Immunofluorescence

Histological analysis and immunocytochemistry were performed (Climer *et al*., 2021) (Murtha et al., 2022). Briefly, cochleae were flushed with 4% PFA and then fixative overnight at 4°C with rotation. Cochleae were then decalcified in 0.1M EDTA for 3 - 5 days at 4°C with rotation. The apical coils of cochleae were then dissected or prepared as 100 μm mid-modiolar sections embedded in the gelatin-agarose solution (Simmons *et al*., 2010; Maison *et al*., 2012). Microdissected cochlear pieces were suspended in 30% sucrose for 30 - 60 min at room temperature with gentle shaking, frozen at -80°C for 30 min, then thawed for 30 min at 37°C. Pieces were washed 3 times in PBS, then blocked in 5% normal horse serum (NHS) for 1 hr at room temperature. Samples were labeled with antibodies to myosin 7A (Proteus Biosciences, USA, No. 25-6790, 1:200), the C-terminal-binding protein 2 (CtBP2, BD Biosciences #612044, 1:200), OCM (Santa Cruz sc-7446, 1:200), GluR2 (Millipore, MAB397, 1:200), and P2X2 (Alomone Labs, Israel, APR-003, 1:400). Primary antibodies were incubated overnight at 37°C. Appropriate Northern Lights (R&D Systems 1:200) and Alexa Fluor (Invitrogen 1:200) conjugated secondary antibodies were incubated for 1 hr at 37°C. Slides were prepared using Vectashield mounting media with DAPI (Vector Laboratories, USA).

### Confocal microscopy

Images were acquired using a Zeiss LSM800 upright confocal laser scanning microscope (Zeiss, Germany). Cohorts of samples were immunostained at the same time and imaged under the same optical conditions to allow for direct comparison. Microdissected cochlear epithelia were imaged using a 10x air objective (N.A. 0.3) at 0.5x zoom to capture entire pieces. Then, higher magnification images were taken using a 40x oil immersion objective (N.A. 1.2). Z-stacks were taken to capture whole hair cells, using Hoechst or DAPI, phalloidin, and Myo7a. Cytocochleograms were constructed from these images by tracing the cochlear spiral and superimposing hash marks using a custom ImageJ Measure Line plugin from Eaton-Peabody^1^. The plugin superimposed frequency correlates on the microdissected spiral image by application of the cochlear frequency map for mice. For cytocochleograms, counting of myosin 7A immunolabeled inner (IHC) and outer (OHC) hair cells was performed in two different 200 µm-long segments of the organ of Corti that corresponded to the 5.66, 8.00, 11.32, 16.00, 22.56, 32 and 45.2 kHz regions from *Ocm*^+/+^ and *Ocm*^-/-^ cochleae. Hair cells were considered absent if the stereociliary bundles and cuticular plates were missing and there was no myosin 7A immunofluorescence. For counts of pre-synaptic ribbons (CtBP2) and ribbon synpases (paired GluR2 and CtBP2), maximum intensity projections of confocal z-stacks either between 16 – 22.6 kHz or between 8 – 12 kHz (apical) of the cochlear spiral were taken. Ribbons were analyzed with Imaris software (v10.2, Oxford Instruments) using the spots feature to identify puncta, and determine their sizes and number. The thresholding criterion for pixel intensity in the CtBP2 channel was set at 30.1 and region threshold set at 191 (for a 8-bit image with a dynamic range of 0–255) depending on the brightness of the projections. The same isointensity criterion was applied to all z-stacks in the same projection. For GCaMP6s and OCM fluorescence intensity, quantification was performed in ImageJ using a region of interest (ROI) from the cytoplasm compared with a similar ROI outside the organ of Corti. After background removal, relative fluorescence intensity was determined. 2-5 cochlea from different animals were analyzed per condition.

### Western Blotting

Cochleae were dissected and harvested in ice-cold PBS. Cochleae were then transferred into a pre-cooled tube (one cochlea per sample) containing 250 µL ice-cold RIPA-lysis buffer containing: 1% Triton X-100, 25 mM Tris, pH 7.4, 150 mM NaCl, 1 mm DTT, 1 mm MgCl_2_, 1 mM phenylmethylsulfonyl fluoride, and 1x protease inhibitor cocktail (Thermo Scientific). A homogenizer (Kimble Pellet Pestle, DWK Life Science) was used to crack up cochleae manually, followed by an additional 30 min incubation in lysis buffer. Cochlear lysates were further sonicated (Fisher Scientific) 3 times x 5 seconds each on ice. Samples were then transferred to a rotating wheel at 4°C for an additional 60 min, lysates were spun down at 4°C, 12000 x g for 5 min to remove cell debris. The supernatants were then transferred to a new tube and heated for 30 minutes at 37°C in 4X SDS-sample buffer (BIO-RAD, 355 mM β-ME added). Samples were then subject to SDS-PAGE and protein was transferred to a PVDF membrane. Membranes were blocked with 2.5% fish gelatin for 1 hr at RT and incubated overnight at 4°C with primary antibody to rabbit P2X2R (1:400, Alomone Labs) and rabbit β-tubulin (1:1000). After washing, membranes were incubated with appropriate HRP-conjugated secondaries (anti-rabbit 1:7,500) for 1 hr at RT. Clarity Max ECL Substrate (BIO-Rad, USA) was used to detect chemiluminescence via a ChemiDoc imaging system. Relative protein expression levels (normalized to β-tubulin) were determined by densitometric analysis using FIJI software.

### Statistical analysis

Statistical analysis was performed using GraphPad Prism 9 and MATLAB software. Statistical comparisons of means were made by t-test or, when normal distribution could not be assumed, the Mann-Whitney U-test. For multiple comparisons, one-way, two-way and three-way ANOVA was followed by unpaired *t*-tests using a pooled variance estimate, with Bonferroni correction unless otherwise stated. A significance level of *p* < 0.05 was used to determine statistical significance. Data are given as mean ± SD. Animals of either sex were randomly assigned to the different experimental groups. No statistical methods were used to define the sample size, which was selected based on previously published similar work from our laboratories. Animals were taken from multiple cages and breeding pairs.

## Results

### OHCs from Ocm^-/-^ mice exhibit normal biophysical profiles and synaptic innervation

At pre-hearing ages, OHCs from *Ocm*^-/-^ mice exhibit normal biophysical profiles (Murtha *et al*., 2022), but exhibit a downregulation of Cav1.3 channels and upregulation of P2X2 receptors (Yang *et al*., 2023). Both Cav1.3 channels and P2X2 receptors play critical roles in shaping the biophysical properties and functional behavior of cochlear hair cells. Therefore, we investigated whether the lack of OCM had any effect on mature OHC biophysical profiles and electromotility. In 1-2-month-old (mo) CBA/CaH mice, we investigated the potassium currents in OHCs while applying a series of hyperpolarizing and depolarizing voltage steps from the holding potential of -84 mV with 10 mV increments (**Figure 1A, B**). The time course and voltage dependence of the outward K^+^ currents in OHCs were comparable between *Ocm*^-/-^ and control OHCs (**Figure 1A-C**). We then tested OHC electromotile responses. Under whole-cell patch clamp conditions, stepping the membrane potential from −84 mV to +40 mV caused OHCs to shorten, as previously described (Marcotti & Kros, 1999; Jeng *et al*., 2020). OHC contraction was not significantly different between the two genotypes (**Figure 1D, E**). Similar to immature OHCs, the lack of OCM does not affect OHC electrophysiology in young adult mice.

**Figure 1.**
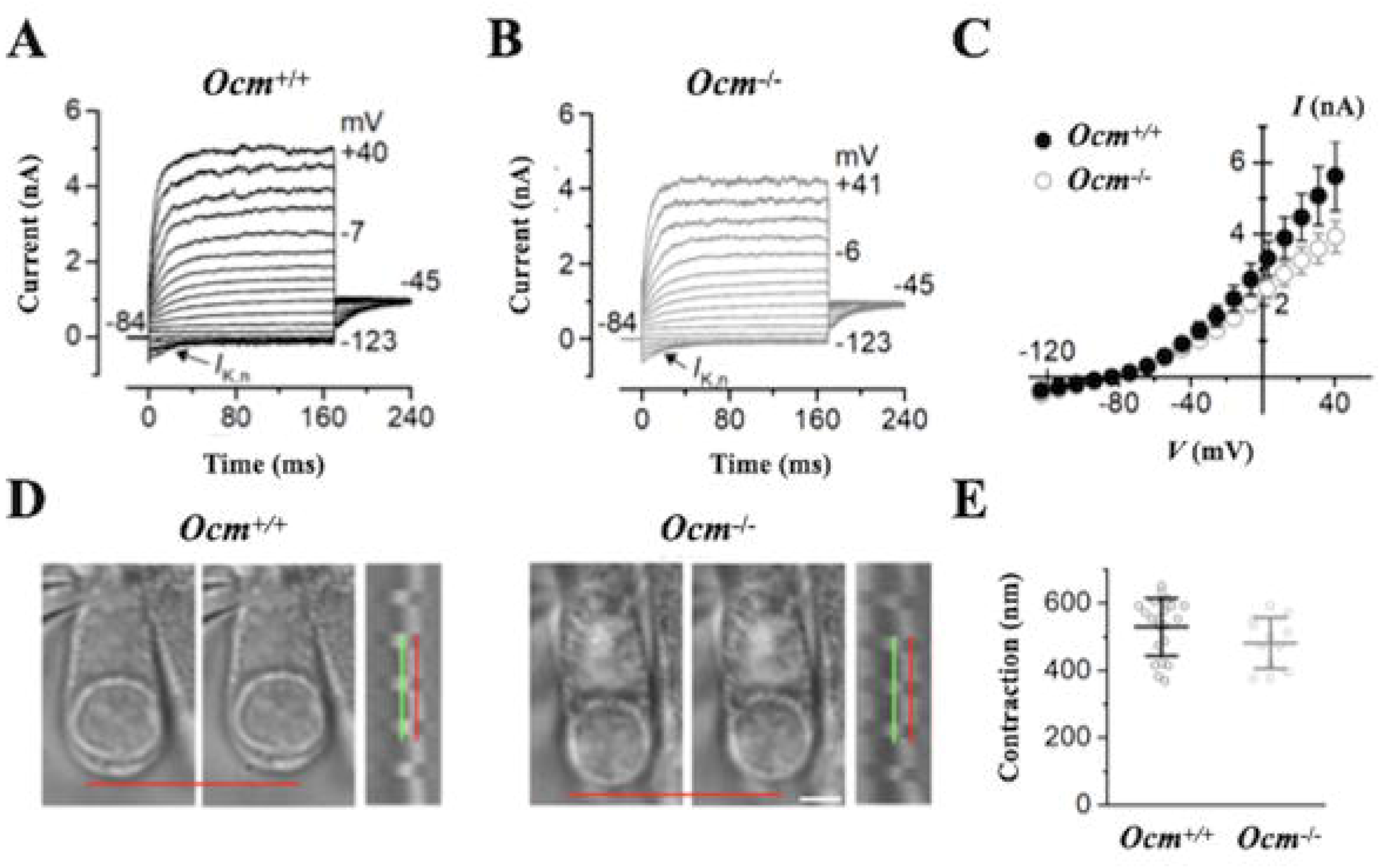
Biophysical properties of OHCs are not affected in early postnatal CBA/CaH *Ocm*^-/-^ mice. K^+^ currents from control OHCs **(A)** and *Ocm*^-/-^ **(B)** 1 - 2 months age CBA/CaH mice. Recordings were performed by applying a series of voltage steps in 10 mV nominal increments from the holding potential of -84 mV. **(C)** Average current-voltage curves from control and *Ocm*^-/-^ OHCs of 1-2 months, n = 4 for each genotype, mean ± SEM. **(D, E)** *Ocm*^-/-^ OHCs showed a comparable contraction distance. n = 7, 21 control OHCs, n = 4, 12 *Ocm*^-/-^ OHCs were measured, mean ± SD with single recordings in open circles, *t*-test.

Hair cells have specialized pre-synaptic ribbons that facilitate neurotransmitter release. In the mature mouse cochlea, each OHC is innervated by two to three type II afferent fibers associated with presynaptic ribbons, which is also about 1/10th of the ribbons present in adult IHCs (Huang *et al*., 2012; Ceriani *et al*., 2019; Johnson *et al*., 2019). During the prehearing period, apical-coil OHCs from *Ocm*^-/-^ mice show increased numbers of ribbons and type II afferent contacts (Yang *et al*., 2023). We investigated whether synapses and ribbons were altered in IHCs and OHCs in the *Ocm*^-/-^ young adult mice. Using CtBP2 and GluR2 immunofluorescence, we focused our study on midcochlear (16 - 22.6 kHz) regions (**Figure 2A-D).** Although both the number of ribbons and synaptic puncta trended higher in *Ocm*^-/-^ mice, they were not statistically different. In IHCs, the mean number of synaptic ribbons was 19.338±1.834 in *Ocm*^-/-^ mice compared to 18.265±0.971 (*p* = 0.4058) in *Ocm*^+/+^ mice (**Figure 2E**). There was also no statistical difference in the mean number of orphan ribbons in *Ocm*^-/-^ IHCs (1.470±1.052) compared to *Ocm*^+/+^ IHCs (1.290±0.563, *p* = 0.974). The mean number of ribbons present in OHCs was 2.330±0.2350 in *Ocm*^-/-^ mice compared to 1.980±0.2750 (*p* = 0.114) in *Ocm*^+/+^ mice (**Figure 2F**). However, ribbon volumes were larger in *Ocm*^-/-^ OHCs (0.142±0.088 µm^3^) compared to *Ocm*^+/+^ OHCs (0.0883±0.040 µm^3^, *p* < 0.0001, *two-way* ANOVA) (**Figure 2G**).

**Figure 2.**
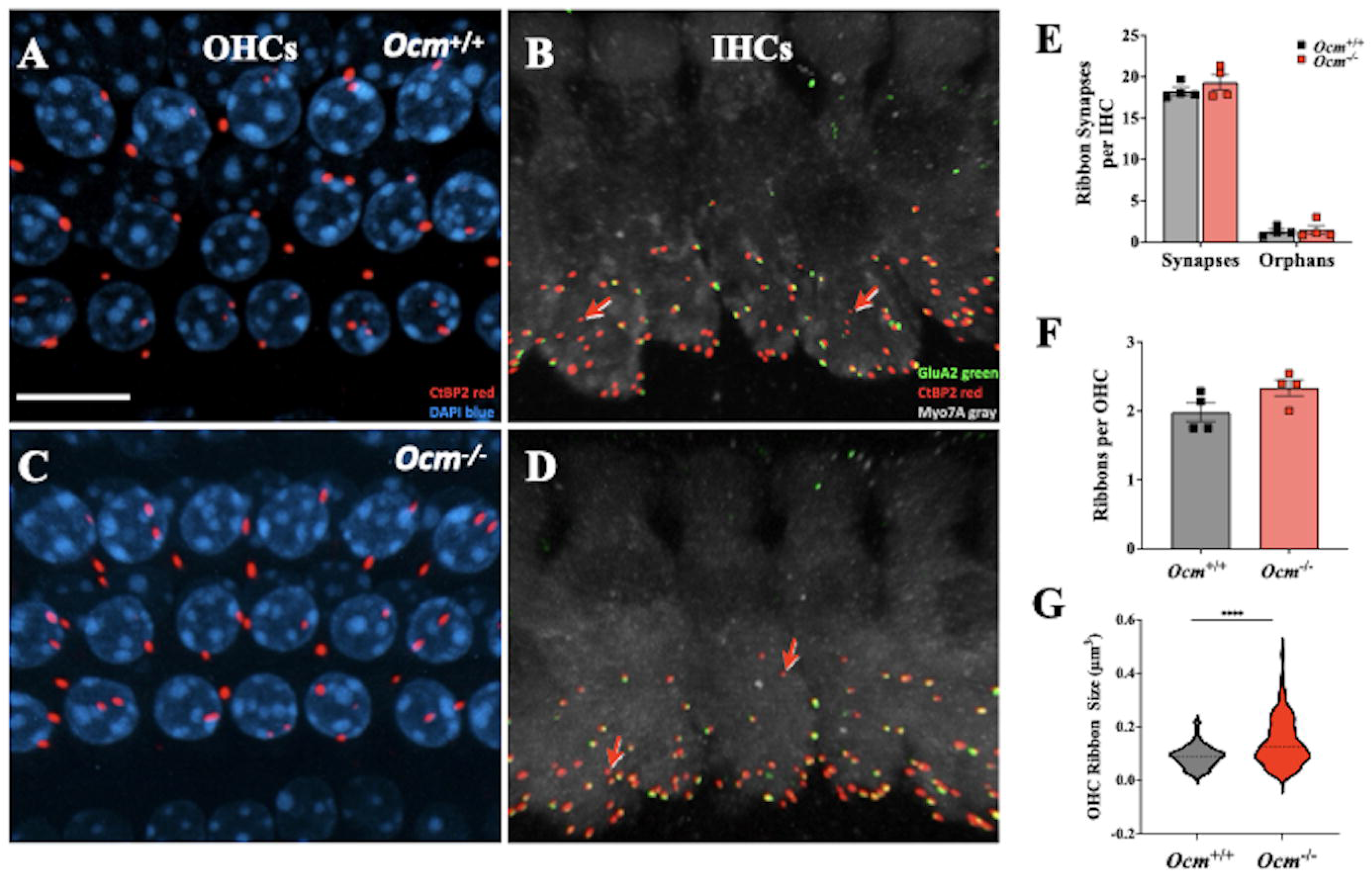
Presynaptic ribbons and postsynaptic glutamate receptors are normal in adult *Ocm*^-/-^ mice. Maximum projections of confocal z-stacks obtained from the 16 kHz region from 1-month old *Ocm^+/+^* (**A, B**) and *Ocm^-/-^* (**C, D**) cochlea. Cochlear whole mounts were immunolabeled for presynaptic ribbons (CtBP2, red), postsynaptic glutamate receptors (GluR2; green), hair cells (myosin VIIA; gray), and nuclei (Hoechst, blue). **A, C** show presynaptic ribbons in the OHC region in *Ocm^+/+^* and *Ocm^-/-^* mice. **B, D** show juxtaposition between CtBP2 and GluR2 synaptic complexes as well as orphaned ribbons (CtBP2 only, red arrows). **E**. The number of glutaminergic ribbon synapses was quantified for IHCs in *Ocm^+/+^* and *Ocm^-/-^* mice. **F**. The number of ribbons (CtBP2 puncta) was quantified for OHCs in *Ocm^+/+^* and *Ocm^-/-^* mice. **G**. The volume of ribbons (CtBP2 puncta) was quantified for OHCs in *Ocm^+/+^* and *Ocm^-/-^* mice.

### Lack of OCM alters temporal coding and increases permanent threshold shifts after noise exposure

We investigated whether the absence of OCM in OHCs increases vulnerability to moderate levels of noise exposure in young adult mice. We used an octave band noise (8-16 kHz) at 95 dB SPL for 2 hrs, designed to cause a large temporary threshold shift (TTS) with minimal or no permanent threshold shift (PTS) (Hirose & Liberman, 2003; Fernandez *et al*., 2010) (**Figure 3A**). To assess cochlear function, we measured auditory brainstem responses (ABRs) and distortion product otoacoustic emission (DPOAE) in anesthetized mice. Since genetic disruption of *Ocm* leads to progressive hearing loss (Climer *et al*., 2021), noise exposure was performed in CBA/CaJ mice that were 4 - 9 weeks, which is well before the appearance of ABR threshold elevation. We compared ABR amplitudes, latencies and thresholds before and again at 6 hrs and 2 wks after noise exposure. Prior to noise exposure, ABR thresholds were similar between wild-type and *Ocm*^-/-^ mutant mice (**Figure 3B-D**). At 6 hrs post-noise exposure, ABR thresholds at 32 kHz showed a significant increase in *Ocm*^+/+^ mice (*p* = 0.0303, *two-way* ANOVA, **Figure 3C**), while in *Ocm*^-/-^ mice thresholds at all three test frequencies were significantly elevated (*p* = 0.0098 for 8 kHz, *p* = 0.0416 for 16 and *p* = 0.0164 for 32kHz, *two-way* ANOVA **Figure 3D**). ABR threshold shifts at 16 kHz were comparable between *Ocm*^+/+^ and *Ocm*^-/-^ mice 6 hrs after noise exposure. However, after 2 wks, thresholds in *Ocm*^-/-^ mice did not recover at 16 kHz in contrast to wild types (*p* = 0.0135 **Figure 3E**).

**Figure 3.**
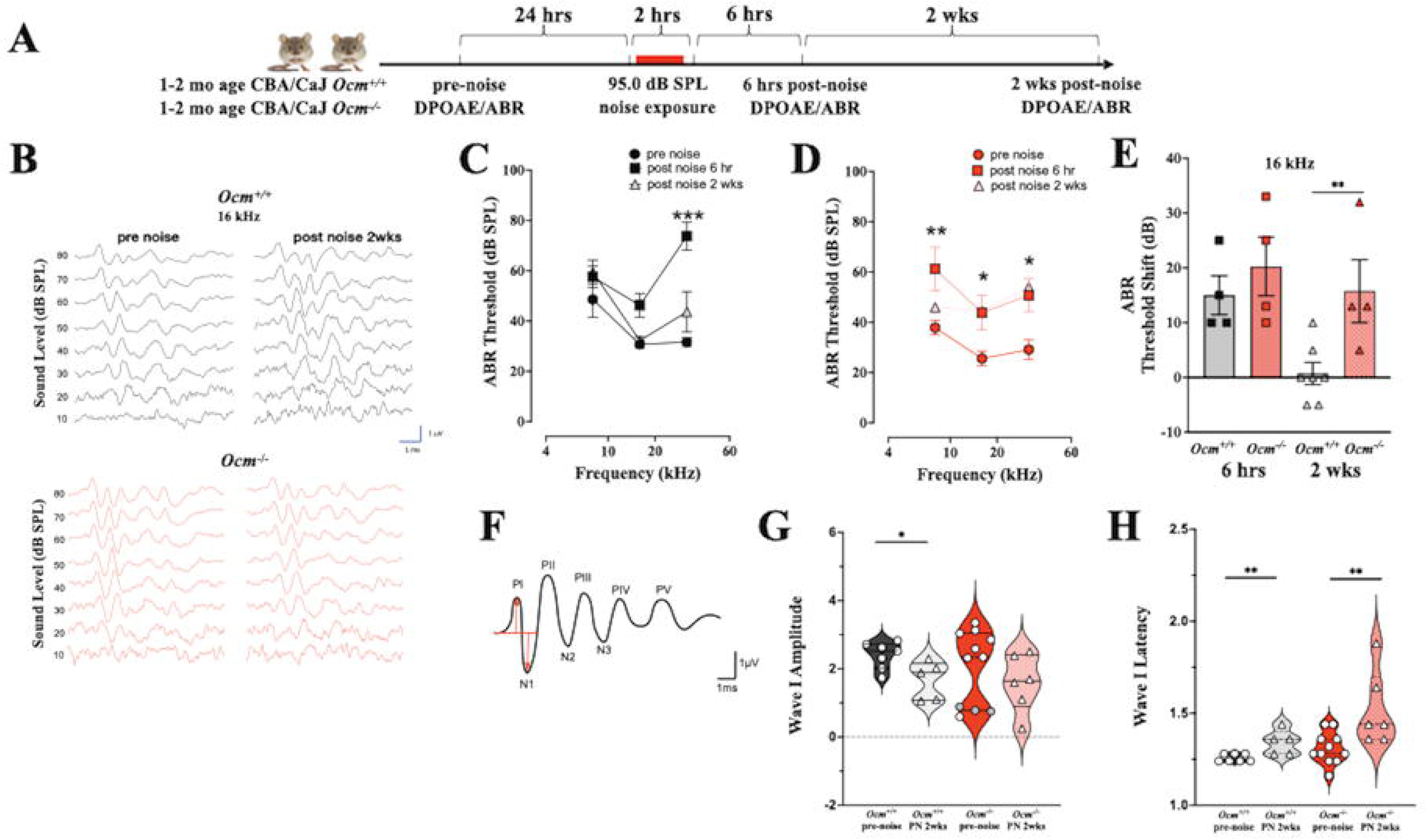
*Ocm^-/-^* mice showed increased ABR threshold shifts after noise exposure. **(A)** Schematic of noise exposure experiment timeline using CBA/CaJ *Ocm^+/+^* or *Ocm^-/-^* mice at 1-2 months of age (mo). DPOAEs and ABRs were tested 24 hrs prior to the noise, followed by 2 hrs 95.0 dB SPL broadband noise exposure. DPOAEs were measured 6 hrs after, and 2 wks after noise exposure. **(B)** Representative ABR waveforms at 16 kHz from CBA/CaJ *Ocm^+/+^* and *Ocm^-/-^* mice before 6 hrs after, and 2 wks after noise exposure. **(C)** ABR thresholds from 1-2 mo CBA/CaJ *Ocm^+/+^* (black/left) and *Ocm^-/-^* (red/right) mice at 8 kHz, 16 kHz, and 32 kHz before, 6 hrs, and 2 wks after noise exposure. n ≥ 4 for each genotype, mean ± SD, *p* < 0.001 for both *Ocm^+/+^* and *Ocm^-/-^* mice between pre-noise and 6 hrs post-noise, *two-way ANOVA*. Significance shown as asterisks (*) indicates significant differences between pre-noise and 6 hrs post noise, no significant differences between pre-noise and 2 wks post noise. **(D)** Average of ABR threshold shifts at 16 kHz, mean ± SD, *p* < 0.05, *two-way ANOVA*. *: *p* < 0.05, **: *p* < 0.01, ***: *p* < 0.001, *Bonferroni* post-test. **(E)** ABR threshold shift at 16 kHz from *Ocm^+/+^* and *Ocm^-/-^* mice before, 6 hrs after, and 2 wks after noise exposure. mean ± SD. **: *p* < 0.01, t-test. **(F)** An idealized ABR waveform showing positive peak (PI – PV) and negative peak (N1 – N3) potentials. The latency and amplitudes of wave I are identified by arrows. **(G, H)** Violin plots analyzing amplitude and latency of ABR waveform potentials wave I (difference between first positive peak potential and first negative peak potential, PI – N1) evoked by a 16 kHz tone at 80 dB SPL. *: *p* < 0.01, t-test. **: *p* < 0.001, *t-*test.

Prior to noise exposure, *Ocm*^-/-^ mice showed larger variability in the amplitude of ABR wave I (N1 – P1 in **Figure 3F**) than wild-type mice. Focusing on 16 kHz at 80 dB SPL, coefficients of variation (CVs) for wave I amplitudes were 53% in *Ocm*^-/-^ mice compared to 17% in wild types. As shown in **Figure 3G**, there was a bimodal distribution of wave I amplitudes in *Ocm*^-/-^ mice prior to noise exposure, with one peak at 2.8 µV (±0.40, N=7, CV=14%) and a second at 0.75 µV (±0.12, N=4, CV=16%). *Ocm*^-/-^ mice had a mean wave I amplitude of 2.058±1.082 µV vs. 2.394±0.3954 µV for *Ocm*^+/+^ mice. In contrast, before exposure, *Ocm*^-/-^ and *Ocm*^+/+^ mice showed similar latencies for wave I (1.3±0.086 ms and 1.26±0.021 ms, respectively, **Figure 3H**).

Two wks after exposure, *Ocm*^-/-^ and wild-type mice showed greater variation in ABR responses (**Figure 3G, H)**. The largest differences between mutant and wild-type mice occurred in their wave I amplitudes. In both *Ocm*^-/-^ and wild-type mice, wave I amplitudes decreased (*Ocm*^-/-^: 1.60±0.84 µV vs wild type: 1.70±0.56 µV), but only significantly so from pre-noise values in the wild-type (*p* = 0.0480) (**Figure 3G**). Again, *Ocm*^-/-^ mice demonstrated higher CV (53%) compared to wild-type mice (33%). As shown in **Figure 3H**, after the noise, *Ocm*^-/-^ mice demonstrated a significant increase in mean latency for wave I (Δ = 1.50±0.20 ms, p = 0.0084, *t test, two-tailed, unpaired,* Mann-Whitney). Wild-type mice had a smaller, but statistically significant increase in wave I latency (Δ = 1.34±0.067 ms, *p* = 0.0189).

DPOAE thresholds were also significantly increased in both *Ocm*^+/+^ and *Ocm*^-/-^ mice 6 hrs after noise exposure (*p* < 0.0001 for *Ocm*^+/+^ and *Ocm*^-/-^ mice, *two-way ANOVA*, **Figure 4A**). After 2 wks, DPOAE thresholds from *Ocm*^+/+^ mice had recovered to pre-noise thresholds, except at 32 kHz (p < 0.0001, *two-way* ANOVA). In contrast, *Ocm*^-/-^ mice still exhibited elevated DPOAE thresholds at all high frequencies (16 kHz, *p* = 0.0015; 22 kHz, *p* = 0.0031; 32 kHz*, p =* 0.0159; 45 kHz, *p* = 0.0202; **Figure 4B**). The mean DPOAE threshold shifts across all frequencies, 6 hrs after noise were comparable between *Ocm*^+/+^ and *Ocm*^-/-^ mice (**Figure 4C**). Two weeks after noise, *Ocm*^-/-^ mice showed significantly larger DPOAE threshold shifts than *Ocm*^+/+^ mice (*p* = 0.0172. *two-way* ANOVA, **Figure 4C**). DPOAE suprathreshold amplitudes at 16 kHz were reduced in both in *Ocm*^+/+^ and *Ocm*^-/-^ mice at 6 hrs post noise (**Figure 4D,E**). However, 2 wks after noise, *Ocm*^-/-^ mice showed lower suprathreshold DPOAE amplitudes, while *Ocm*^+/+^ mice recovered. The near complete recovery from exposure in wild types suggests that OCM may be involved in protection from PTS after moderate noise exposures.

**Figure 4.**
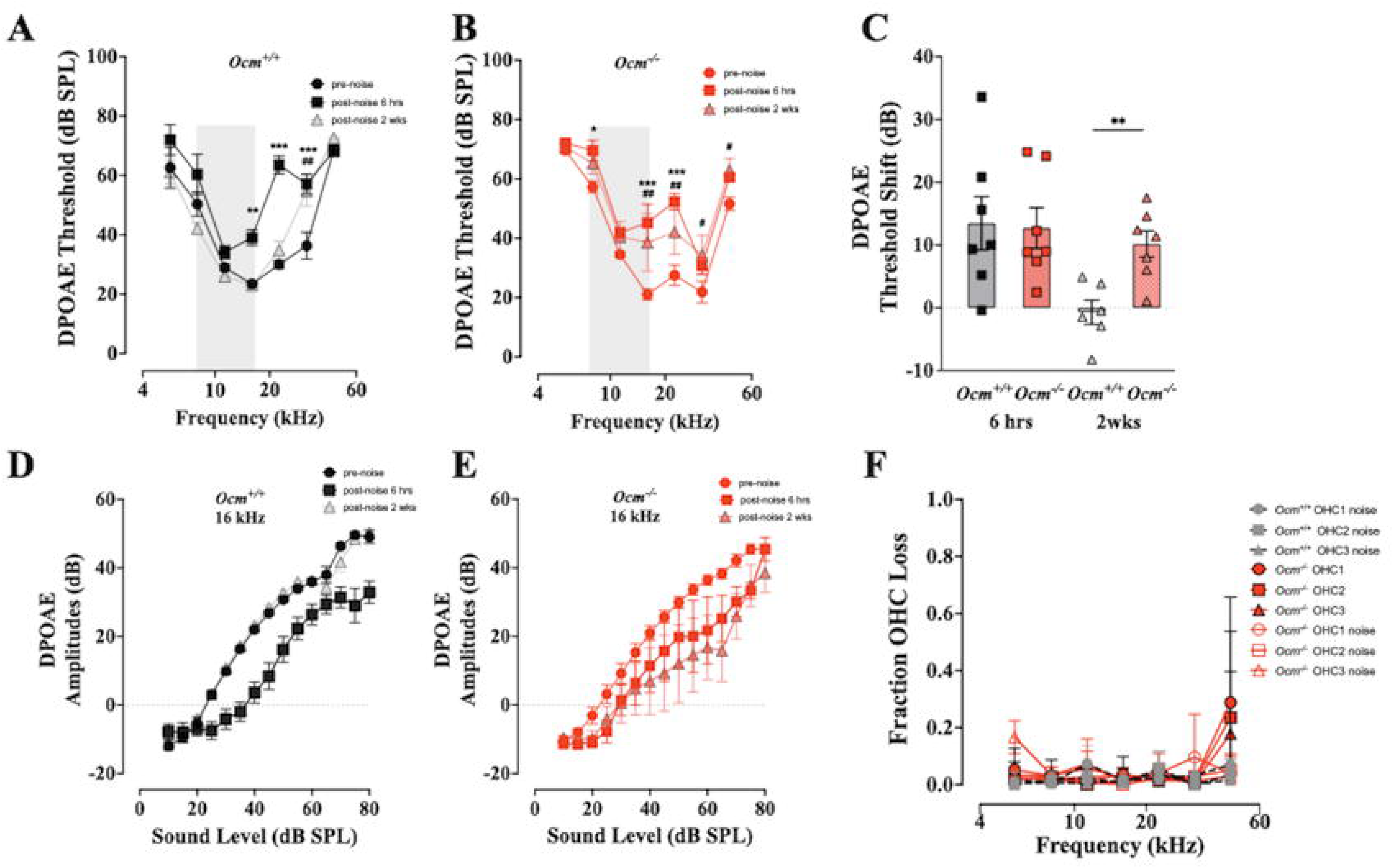
*Ocm^-/-^* mice showed increased DPOAE threshold shifts after noise exposure. **(A, B)** DPOAE thresholds from 1-2 mo CBA/CaJ *Ocm^+/+^* and *Ocm^-/-^* mice before, 6 hrs after, and 2 wks after noise exposure. n ≥ 7 for each genotype. Mean ± SD, *p* < 0.001 for both *Ocm^+/+^* and *Ocm^-/-^* mice between pre-noise and 6 hrs post-noise, and *Ocm^-/-^* mice between pre-noise and 2 wks post-noise *two-way* ANOVA. Significance shown as asterisks (*) means the comparison between pre-noise and 6 hrs post-noise, and the number signs (#) represents the comparison between pre-noise and 2 wks post-noise. */#: *p* < 0.05, **/##: *p* < 0.01, ***: *p* < 0.001, post-test from *two-way* ANOVA. Specifically, *Ocm*^+/+^ mice showed significantly increased DPOAE thresholds at 16 kHz (p < 0.01, *Bonferroni* post-test), 22 and 32 kHz (*p* < 0.0001, *Bonferroni* post-test), while *Ocm*^-/-^ mice showed significantly higher DPOAE thresholds at 8 kHz (p < 0.05, post-test from *two-way* ANOVA), 16 and 22 kHz (p < 0.0001, post-test from *two-way* ANOVA). **(C)** Average of total DPOAE threshold shifts from all frequencies, mean ± SD, *p* < 0.05, *two-way* ANOVA. *: *p* < 0.05, *Bonferroni* post-test. (**D, E)** Mean amplitude versus level functions (±SEM) at 16 kHz for DOAEs in *Ocm^+/+^* and *Ocm^-/-^* mice. (**F**) Mean OHC loss at seven frequency regions (5.6. 8.0, 11.3, 16.0, 22.6, 32.0, and 45.2 kHz)

Permanent threshold shifts after noise are often associated with damaged or missing OHCs, whereas TTS is not (Liberman & Mulroy, 1982). We assessed OHC loss 2 wks after noise exposure. As expected, 95 dB SPL noise produced no significant OHC loss in *Ocm*^+/+^ mice (**Figure 4F**). Similarly, the *Ocm*^-/-^ mice demonstrated no evidence of OHC loss in either control or exposed ears except at 45 kHz, the highest frequency evaluated. Although *Ocm*^-/-^ mice showed significant ABR (**Figure 3E**) and DPOAE (**Figure 4C**) threshold shifts across frequency regions at 2 wks post exposure, there was little hair cell loss associated with this PTS except for 45 kHz (**Figure 4F**).

### Lack of OCM increases threshold shifts following prolonged noise exposure

Noise exposure dramatically increases the levels of intracellular free Ca^2+^ in OHCs (Fridberger *et al*.; Zuo *et al*., 2008). To investigate Ca^2+^ activity, we expressed a tissue-specific Ca^2+^ sensor (*Atoh1*-GCaMP6s) in *Ocm*^+/+^ and *Ocm*^-/-^ mice as previously described (Yang *et al*., 2023). Since the loss of OCM also causes increased levels of intracellular free Ca^2+^ in postnatal mice (Murtha *et al*., 2022), we hypothesized that the early progressive hearing loss observed in adult Ocm^-/-^ mice is partly due to prolonged Ca^2+^ overload in OHCs. We compared *Ocm^+/+^* and *Ocm^-/-^* in their susceptibility to prolonged (9-hr) broadband noise exposure at 95 dB SPL. Cochlear thresholds were measured 1 hour following noise exposure (**Figure 5A**). As a Ca^2+^ sensor, GCaMP6s contains a calmodulin domain that might also disrupt calcium homeostasis over time. However, at 3 - 4 wks of age, neither the endogenous expression of GCaMP6s nor the absence of OCM affects DPOAE thresholds at any frequency (*p* = 0.6928 for *Ocm*^+/+^ vs. *Ocm*^-/-^, *p* = 0.2887 for GCaMP6s(-) [*cre*-] vs. GCaMP6s(+) [*cre*+], *three-way ANOVA*, **Figure 5B**). We measured ABR thresholds only at 16 kHz, and before exposure there were no significant differences among GCaMP6s(-) and GCaMP6s(+), *Ocm^+/+^* and *Ocm^-/-^* mice (*p* = 0.4744, *two-way ANOVA*, **Figure 5C**). After 9 hrs of exposure at 95 dB SPL, *Ocm^-/-^* mice showed a larger ABR threshold shifts (*p* = 0.0178, *t*-test, **Figure 5D**). In *Ocm^+/+^* mice, DPOAE thresholds were significantly elevated only at 22 kHz (*p* = 0.0430, *two-way ANOVA*, **Figure 5E**). However, in *Ocm^-/-^* mice, DPOAE thresholds were elevated at 11 (*p* = 0.0010), 16 (*p* < 0.0010), and 22 kHz frequencies (*p* = 0.0027, **Figure 5F**). The mean DPOAE threshold shift across all frequencies was significantly higher in *Ocm^-/-^* mice compared to wild types (*p* = 0.04, *t*-test, **Figure 5G**). Thus, GCaMP6s(+);*Ocm*^-/-^ mice demonstrated similar vulnerability to prolonged broad-band noise as CBA/CaJ *Ocm*^-/-^ mice did for shorter duration octave-band noise exposure.

**Figure 5.**
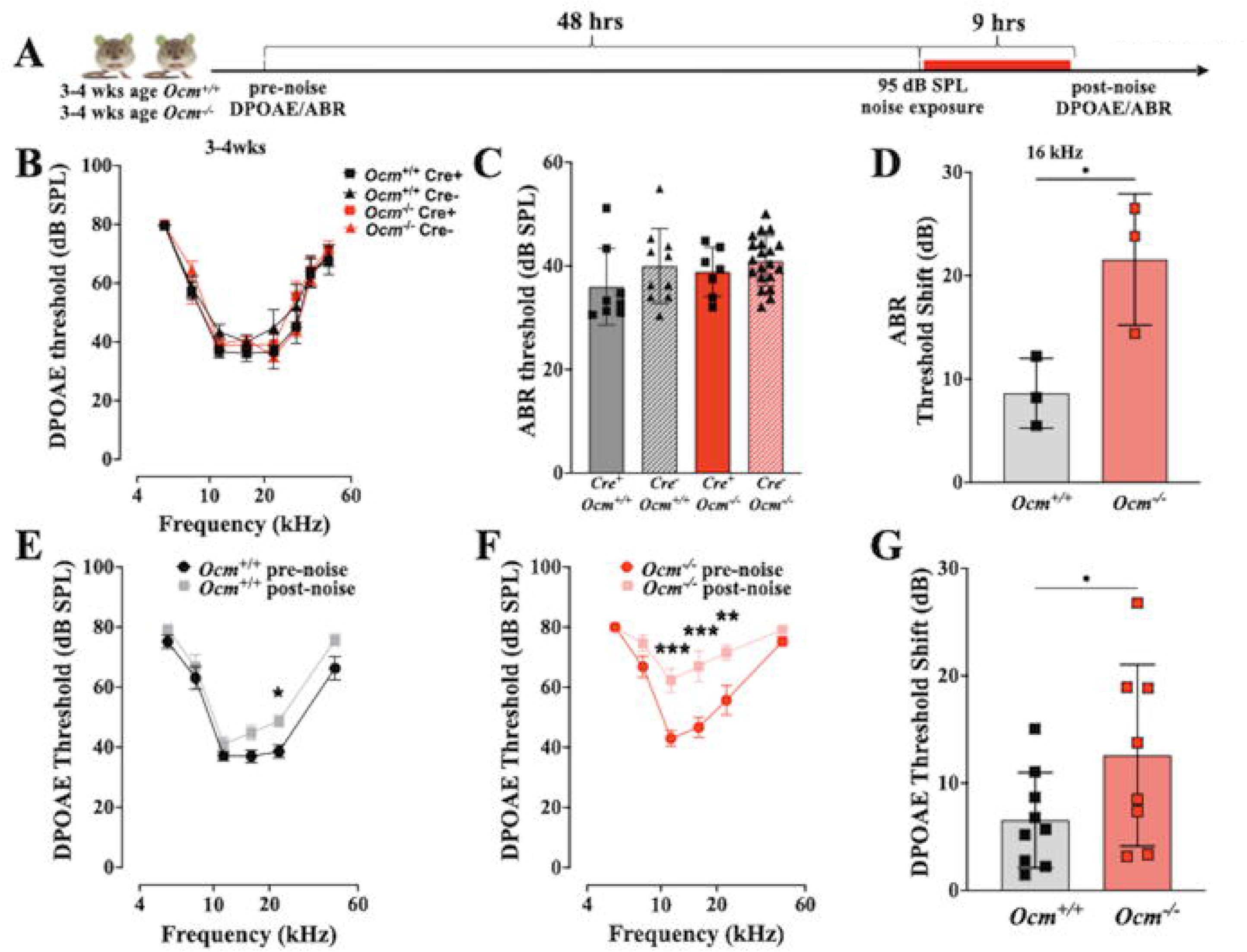
*Ocm^-/-^* mice showed increased susceptibility after prolonged noise exposure. **(A)** Schematic of noise exposure experiment timeline using *Ocm^+/+^* or *Ocm^-/-^* mice with endogenous expression of GCaMP6s at 3 - 4 weeks (wks) of age. ABRs and DPOAEs were tested 48 hours (hrs) prior to noise, followed by a 9 hrs 95 dB SPL broadband noise exposure. DPOAEs were then measured immediately after noise exposure. **(B)** DPOAEs were measured from GCaMP6s *Ocm^-/-^* mice and age-matched GCaMP6s *Ocm^+/+^* mice at 3 - 4 wks, n = 10 *Ocm*^+/+^ GCaMP6s-mice, n = 8; *Ocm*^+/+^ GCaMP6s+ mice, n = 20; *Ocm^-/-^* GCaMP6s-mice, and n = 7 *Ocm^-/-^* GCaMP6s+ mice. **(C)** ABR thresholds at 16 KHz, Cre+ means the endogenous expression of GCaMP6s, mean ± SD. **(D)** ABR threshold shift at 16 kHz from 3 - 4 wks old *Ocm^+/+^* and *Ocm^-/-^* mice before and after noise exposure. mean ± SD. **: *p* < 0.01, t-test. **(E - F)** DPOAEs from 3 - 4 wks old *Ocm^+/+^* and *Ocm^-/-^* mice before and after noise exposure. n = 6 for each genotype. *p* < 0.01 for *Ocm^+/+^* and *Ocm^-/-^* mice, *two-way* ANOVA. Significance shown as asterisks, *: *p* < 0.05, **: *p* < 0.01, ***: *p* < 0.001, *Bonferroni* post-test. **(G)** The average of total DPOAE threshold shifts was measured from all frequencies, mean ± SD. *: *p* < 0.05; *t*-test.

### Prolonged noise exposure induces higher GCaMP6s signaling in Ocm^-/-^ mice

Both *Ocm*^+/+^ and *Ocm*^−/−^ mice showed endogenous GCaMP6s fluorescence in the cochlea under confocal microscopy (**Figure 6A**). Before noise exposure, *Ocm*^−/−^ OHCs had significantly higher baseline fluorescence intensity than *Ocm*^+/+^ OHCs. As shown in **Figure 6B**, relative fluorescence intensity for OHCs in *Ocm*^+/+^ mice was 45.900±12.980, while in *Ocm*^−/−^ mice it was 54.430±21.380 (*p* = 0.0280, *Kruskal-Wallis test*). However, immediately following noise exposure, there was a 31.18% increase in *Ocm*^−/−^ OHCs fluorescence (71.400±40.700, *p* = 0.0060, *Kruskal-Wallis test*), while little or no change (-4.9%) in *Ocm*^+/+^ OHCs (43.640±14.070, *p* > 0.9990, *Kruskal-Wallis test*). The increased fluorescence following noise in *Ocm*^−/−^ OHCs (**Figure 6B**), may indicate a higher basal level of intracellular Ca^2+^ signaling due to the lack of OCM as previously suggested (Murtha *et al*., 2022; Yang *et al*., 2023). Furthermore, IHCs in *Ocm*^−/−^ mice also exhibited higher fluorescence both before and after noise compared to wild types (**Figure 6C**). Before noise exposure, IHCs from *Ocm*^+/+^ and *Ocm*^−/−^ mice had relative fluorescent intensities of 88.010±61.740 and 134.800±50.090 (*p* = 0.0003), respectively. After noise, there was a modest increase (10.96%) in fluorescent intensities in *Ocm*^+/+^ IHCs (97.660±50.650, *p* = 0.8339, *Kruskal-Wallis test*). This constrasted with IHCs from *Ocm*^−/−^ mice that showed a 23.66% increase after noise (166.700±57.510, *p* = 0.0276, *Kruskal-Wallis test*).

**Figure 6.**
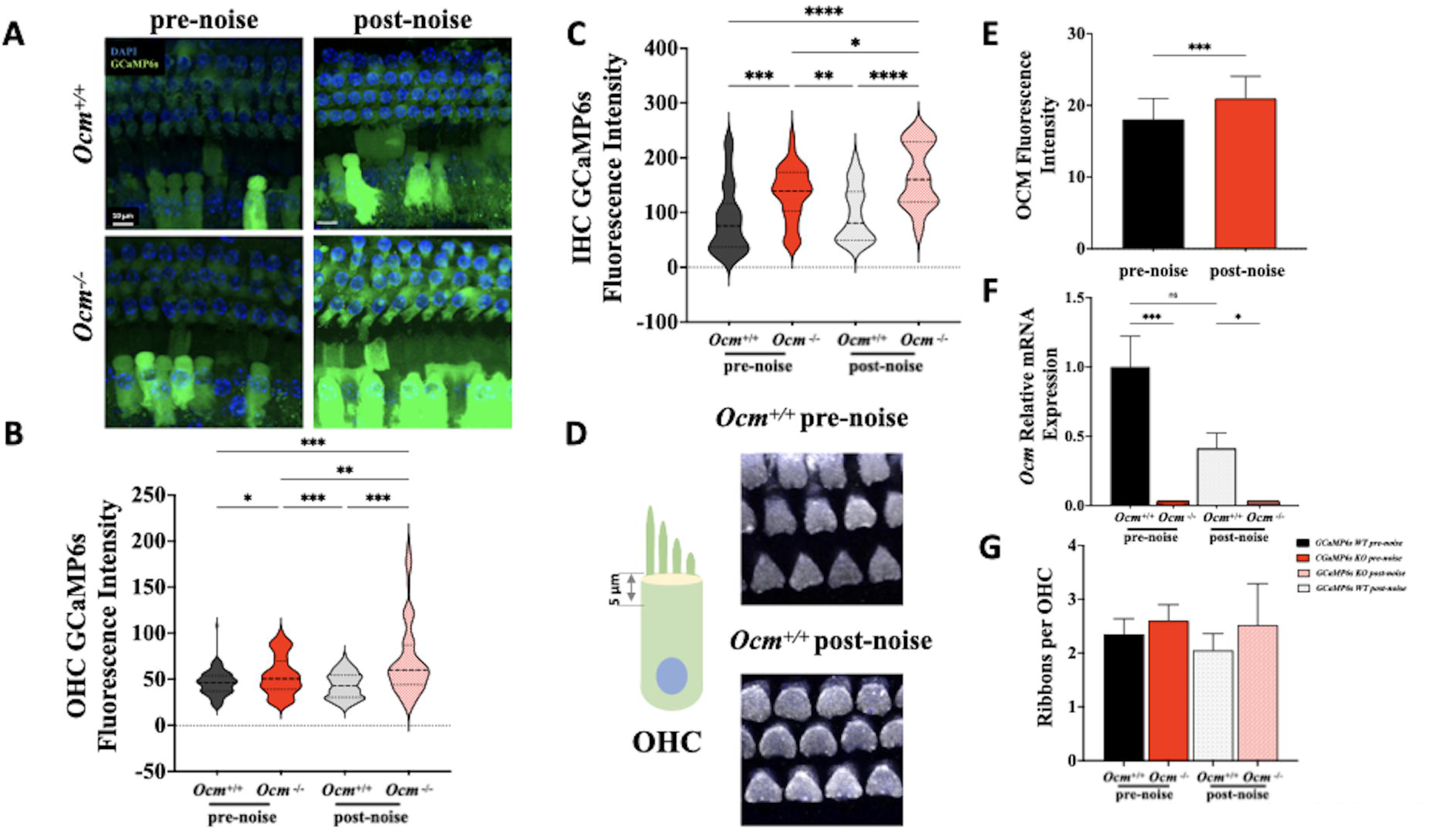
Prolonged noise exposure alters GCaMP6s signaling and OCM expression. **(A**) Maximum intensity projections of confocal z-stack images taken from GCaMP6s+ *Ocm*^+/+^ and *Ocm*^-/-^ apical-spirals before and after prolonged noise exposure. GCaMP6s+ mice showed endogenous green fluorescence (green) in cochlear with the organ of Corti, but most intensely in hair cells. Both OHCs and IHCs from GCaMP6s+ *Ocm*^-/-^ mice showed higher endogenous green fluorescence after noise exposure. OHC (**B)** and IHC **(C)** relative fluorescence intensity were quantified per genotype before and after noise**. (D)**. In GCaMP6s+ *Ocm*^+/+^ mice, OCM (gray) labeling was present at the cuticular plate. There was increased labeling intensity after noise. **(E)** OCM relative fluorescence intensity was quantified before and after noise. **(F)** The relative mRNA expression of Ocm showed no significant difference before and after noise in *Ocm*^+/+^ mice. **(G)** The number of ribbons (CtBP2 puncta) was assessed for OHCs in *Ocm^+/+^* and *Ocm^-/-^* mice.

### Changes in Ocm expression and synaptic ribbons after prolonged noise exposure

In wild-type mice, prevous studies suggest that OCM localizes to the basolateral membrane and the region just below the cuticular plate (Simmons *et al*., 2010). In the present study, prolonged noise increased the fluorescent labeling of OCM at the cuticular plate (**Figure 6D**): 18.060±2.0280 vs. 20.950±3.1150 (*p < 0.0010, t-test* unpaired) (**Figure 6E**). However, there was no significant difference in *Ocm* mRNA expression before and after noise (**Figure 6F**), suggesting that the changes seen in immunofluorescence might represent a change in OCM localization.

We also investigated the response of OHC ribbons immediately following prolonged noise exposure in apical regions of the cochlea (**Figure 6G**). A blinded observer counted CtBP2 immunopuncta from the perinuclear region to the synaptic pole in maximum intensity projections. We found no significant effects of noise exposure or genotype. Before exposure, the average number of CtBP2 puncta per OHC was 2.350±0.2887 in *Ocm*^+/+^ vs. 2.600±0.3000 in *Ocm*^-/-^ mice. Immediately following exposure, ribbon counts were 2.050±0.3109 in *Ocm*^+/+^ and 2.520±0.7727 (*p* > 0.9999) in *Ocm*^-/-^ mice.

### ATP-induced Ca^2+^ transients in mature OHCs are higher in Ocm^-/-^ mice and after prolonged noise exposure

To assess Ca^2+^ activity in *Ocm^+/+^* and *Ocm^-/-^* mice with GCaMP6s expression, apical cochleae from 3 - 4 wk old mice were dissected (**Figure 7A**) and immediately transferred to the microscope chamber. ATP (100 μM) was extracellularly applied to induce Ca^2+^ transients (**Figure 7B**), which were measured as fractional changes of GCaMP6s fluorescence (ΔF/F_0_) (**Figure 7C** gray line). OHCs from *Ocm^-/-^* mice showed significantly larger ATP-induced Ca^2+^ transients compared to *Ocm^+/+^* littermate (*p* = 0.0015, *post hoc* test from *two-way ANOVA*, **Figure 7D**).

**Figure 7.**
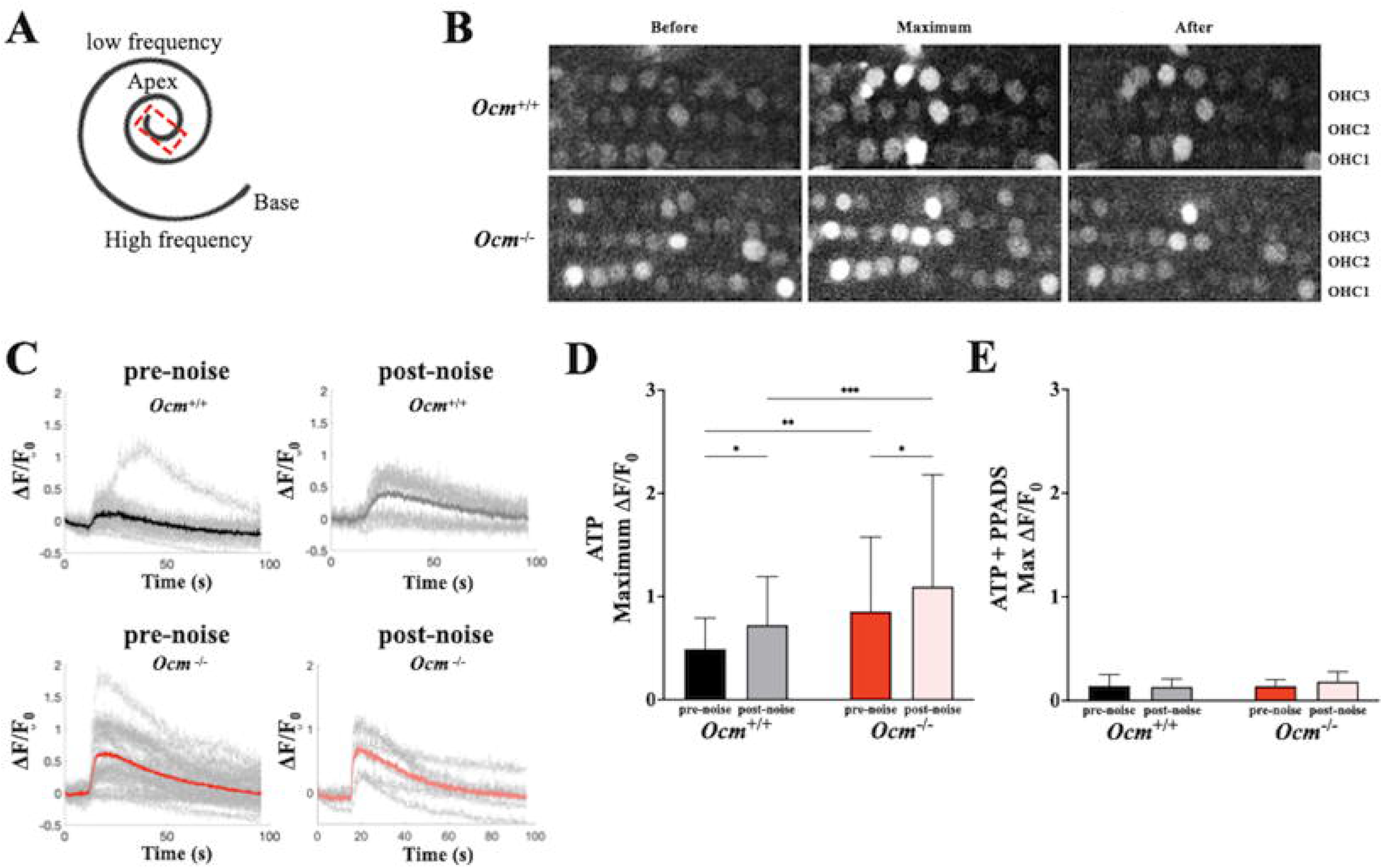
The absence of OCM alters ATP-induced Ca^2+^ transient in OHCs similar to noise. **(A)** Cochleae were taken from 3 - 4 wks old mice with endogenous expression of GCaMP6s. Apical cochlear spirals were immediately dissected and tested. The dashed box shows the region for the Ca^2+^ transient imaging. **(B)** Representative images from pre-noise *Ocm^+/+^* and *Ocm^-/-^* mice showing Ca^2+^ transients in OHCs induced by 100 μM extracellular ATP. GCaMP6s fluorescence before ATP superfusion (top), at peak response (middle), and recovery stage (bottom) are shown. **(C)** Representative traces of fluorescence fractional change (ΔF/F_0_) in OHCs from *Ocm^+/+^* and *Ocm^-/-^* OHCs pre- or post-noise exposure (light gray line), average ΔF/F_0_ shown as bold lines. **(D, E)** Maximum ΔF/F_0_ in OHCs from *Ocm^+/+^* and *Ocm^-/-^* mice pre- or post-noise exposure induced by 100 μM ATP, or with the presence of PPADS (100 μM). For the maximum ΔF/F_0_ of ATP-induced Ca^2+^ transients, n ≥ 4 for each genotype under each treatment, 73 and 93 OHCs from pre-noise *Ocm^+/+^* and *Ocm^-/-^* mice, 83 and 49 OHCs from post-noise *Ocm^+/+^* and *Ocm^-/-^* mice were measured. *p* < 0.001, *two-way* ANOVA. For the PPADS experiment, n = 2 for each genotype under each treatment, 22 and 29 OHCs from pre-noise *Ocm^+/+^* and *Ocm^-/-^* mice, 15 and 13 OHCs from post-noise *Ocm^+/+^* and *Ocm^-/-^* mice were measured. *: *p* < 0.05; **: *p* < 0.01; ***: *p* < 0.001, post-test from *two-way* ANOVA.

We investigated whether the higher susceptibility of *Ocm^-/-^* mice to the prolonged (9 hours) noise exposure was a result of increased Ca^2+^ signaling in OHCs (Murtha et al., 2022; Yang et al., 2023). We found prolonged noise stimulation caused a further increase in maximum ΔF/F_0_ in OHCs from both genotypes (*p* = 0.0030, *two-way ANOVA*, **Figure 7D**). However, OHCs from *Ocm^-/-^* mice exhibited a higher maximum ΔF/F_0_ signal after noise compared to *Ocm^+/+^* mice (*p* = 0.0008, *post hoc* test from *two-way* ANOVA. **Figure 7D**). The maximum ΔF/F_0_ signals between pre-noise OHCs in *Ocm^-/-^* mice and post-noise OHCs in *Ocm^+/+^* mice were similar (*p* = 0.0015, *post hoc* test from *two-way ANOVA*, **Figure 7D**).

To test whether the ATP and noise-induced Ca^2+^ responses in OHCs were mediated by purinergic receptors, we used the non-selective purinergic receptor antagonist PPADS (100 μM). Extracellular application of PPADS nearly abolished ATP-induced Ca^2+^ transients in OHCs and eliminated the noise-induced difference in the maximum ΔF/F_0_ between *Ocm^+/+^* and *Ocm^-/-^* (*p* = 0.2703, *two-way ANOVA*, **Figure 7E)**. These data indicate that the lack of OCM increases ATP-induced Ca^2+^ transients in mature OHCs, which mimic the effects caused by long noise exposure.

### Lack of OCM and prolonged noise stimulation upregulate P2X2 receptors

We tested whether the larger Ca^2+^ responses in adult *Ocm^-/-^* mice were due to the increased expression of P2X2 receptors, which are upregulated during prehearing stages (Yang *et al*., 2023). In apical turn wholemounts, P2X2 receptors were found in the organ of Corti of both *Ocm^+/+^* and *Ocm^-/-^* mice. Before noise exposure, *Ocm^-/-^* but not *Ocm^+/+^* mice showed strong P2X2 receptor labeling in OHCs, especially near the cuticular plate (**Figure 8A**, red arrow). After prolonged noise exposures, *Ocm^+/+^* OHCs showed stronger P2X2 receptor immunolabeling in the stereocila bundles compared to pre-noise *Ocm^+/+^* mice (**Figure 8B**, red arrow). There were no obvious changes between pre- and post-noise *Ocm^-/-^* OHCs.

**Figure 8.**
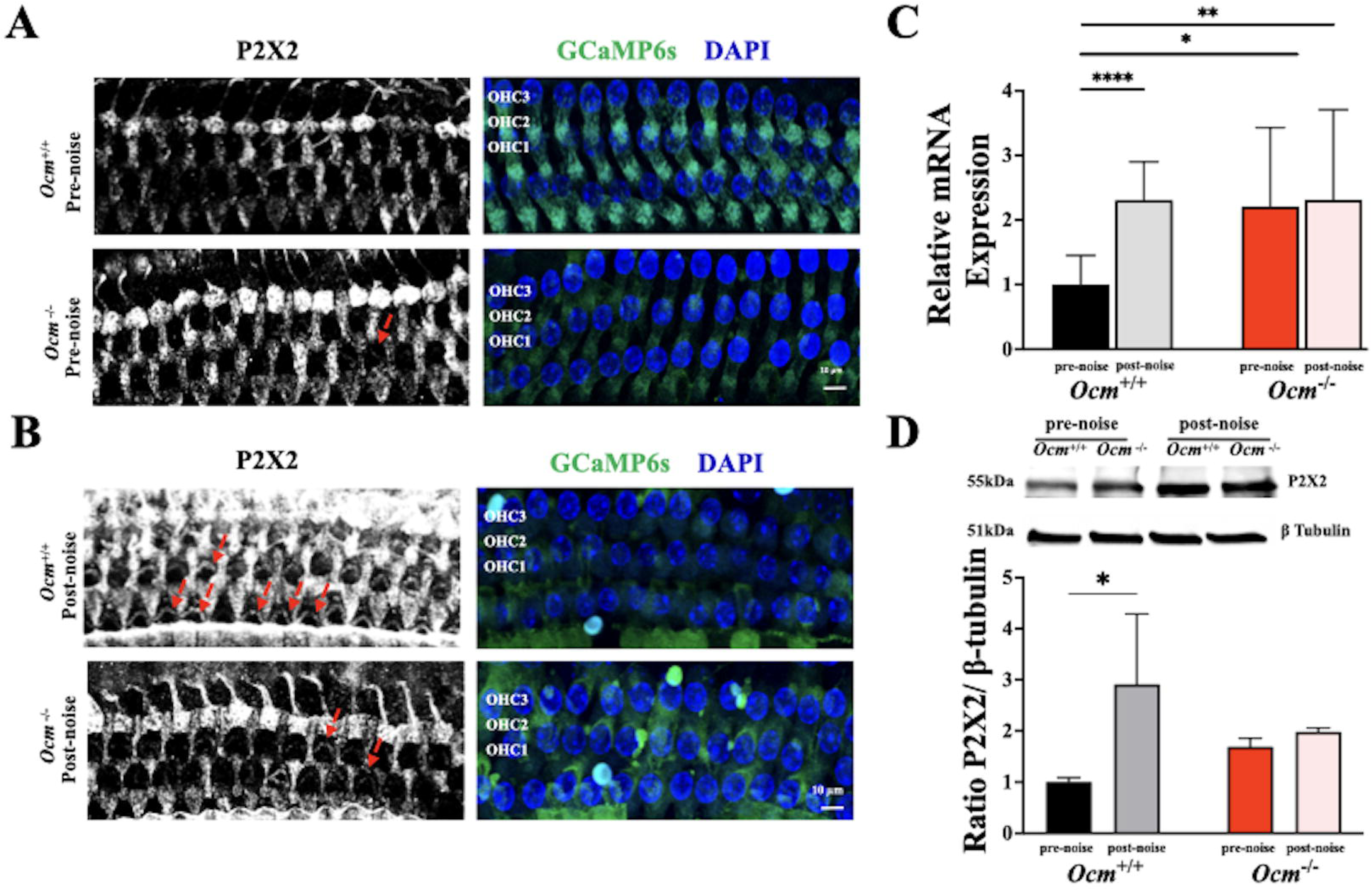
P2X purinoceptor 2 (P2X2) expression upregulated in *Ocm^+/+^* but not *Ocm^-/-^* mice after noise. **(A, B)** Maximum intensity projections of P2X2 immunolabeling on three rows of OHCs harvested from pre- and post-noise *Ocm^+/+^* and *Ocm^-/-^* mice. P2X2 (white), GCaMP6s (green), and DAPI (blue) are shown. Red arrows point to the OHC stereocilia. **(C)** qRT-PCR results of *P2RX2* relative expression, cochleae are taken from 3 - 4 wks *Ocm^+/+^* and *Ocm^-/-^* mice with or without noise exposure. Total mRNA was extracted immediately after dissection. n ≥ 3 for each genotype under each experimental condition. All values were normalized to pre-noise *Ocm^+/+^*. mean ± SD, *p* < 0.05, *t test, two-tailed, unpaired,* Mann-Whitney. **(D)** Representative western blot for P2X2 protein expression levels detected in single cochlea derived from pre- or post-noise *Ocm^+/+^* and *Ocm^-/-^* mice at 3 - 4 wks. n = 3 for each genotype under each experiment condition, β-tubulin (loading control) was used for normalization, all values were then normalized to pre-noise *Ocm^+/+^*. *p* < 0.05 *two-way* ANOVA, ns: not significant, *: *p* < 0.05; **: *p* < 0.01, *Bonferroni* post-test.

Without noise exposure, the cochlear expression of *P2RX2* mRNA in 3-4 wk old *Ocm^-/-^* mice was significantly higher than that in *Ocm^+/+^* mice (*p* = 0.0036, **Figure 8C**). After prolonged noise exposure, *P2RX2* expression was significantly upregulated in the *Ocm^+/+^* cochlea (*p* = 0.0054) but not in *Ocm^-/-^* cochlea (*p* = 0.3472 **Figure 8C**). There was no significant difference in *P2RX2* expression between post-noise *Ocm^+/+^* and *Ocm^-/-^* OHCs (*p* = 0.9933, *post hoc test two-way ANOVA*, **Figure 8C**). Western blot experiments also showed that prolonged noise stimulation significantly upregulated the expression of P2X2 receptors in *Ocm^+/+^* mice (*p* = 0.0395, *post hoc* test from two-way ANOVA, **Figure 8D**). The expression of P2X2 receptor in pre-noise *Ocm^-/-^* mice showed no significant difference (*p* = 0.9523, *post hoc* test from two-way ANOVA, **Figure 8D**). These data suggest that the absence of OCM leads to an upregulation of P2X2 receptors that does not change after prolonged noise exposure.

## Discussion

In the present study, we show that OHCs lacking the Ca^2+^ buffering protein OCM, have increased vulnerability to noise exposure and higher expression of purinergic receptors at young adult ages. Short-duration noise exposure (95 dB SPL, 2 hrs), caused similar acute shifts in the ABR and DPOAE thresholds in *Ocm* control and mutant mice 6 hrs post exposure, but permanent shifts only in *Ocm^-/-^* mice after 2 wks recovery. Moderate noise exposure also increased ABR wave I latencies but had no effect on amplitudes in *Ocm^-/-^* mice, and caused more variability in these neural responses. Prolonged noise exposures (95 dB SPL, 9 hrs) produced larger threshold shifts and increased ATP-induced Ca^2+^ signaling in *Ocm^-/-^* OHCs compared to *Ocm*^+/+^ controls and increased expression of purinergic P2X2 receptors in the cochlea of *Ocm*^+/+^ but not in *Ocm*^-/-^ mice. Upregulated ATP-induced Ca^2+^ responses and higher levels of P2X2 in mature *Ocm^-/-^* OHCs suggest they may have a higher Ca^2+^ load, as previously shown during pre-hearing stages of development (Murtha *et al*., 2022; Yang *et al*., 2023). Additionally, the *Ocm*^-/-^ mice showed modified Ca^2+^ responses to noise in IHCs. All these data confirm that OCM plays a key role in shaping Ca^2+^ dynamics in OHCs that alter cochlear function. The absence of OCM in OHCs causes the cochlea to behave similarly to those in a noise-exposed ear.

Although ABR and DPOAE thresholds are sensitive to hair cell damage, they are relatively insensitive to the loss of cochlear synapses or neural pathology that follows either noise or aging (Kujawa & Liberman, 2009; Stamper & Johnson, 2015; Liberman & Kujawa, 2017). In the present study, *Ocm^-/-^* and wildtype mice had similar threshold shifts 6 hrs after noise exposure, but ABR and DPOAE thresholds never recovered in *Ocm^-/-^* mice as they did in wild-type mice, suggesting that noise caused significant sensory cell damage or loss in the absence of OCM. Since pre-noise ABR and DPOAE thresholds were similar between the two genotypes, we investigated whether the underlying neural responses differed. ABRs are summed, electrical potentials representing the synchronous activity of auditory nerve fibers contacting IHCs (Buchwald & Huang, 1975; Chen *et al*., 2006). Several studies have demonstrated that temporary threshold shifts can lead to permanent loss of IHC synapses with little effect on the recovered ABR and DPOAE thresholds (Lavinsky *et al*., 2015; Mehraei *et al*., 2016; Liberman & Kujawa, 2017). Unlike wild-type mice, the *Ocm^-/-^* mice had highly variable wave I amplitudes, and longer wave I latencies before and after noise exposure. One cause of altered ABR amplitudes is loss of IHC synapses. However in the present study, there was no decrease in synaptic counts in *Ocm^-/-^* mice prior to noise exposure. Reduced amplitudes and increased latencies can also arise when there is damage to the high-frequency region, because lower frequency regions produce responses with longer latencies. At least in unexposed mice, DPOAE thresholds suggest that the OHCs in high-frequency regions are normal, but the highest test frequency was 45 kHz and the extreme base of the mouse cochlea is tuned to frequencies almost an octave higher.

Studies of presynaptic ribbons in OHCs after noise exposure have reported an increase in number and size (Wood *et al*., 2021) when evaluated 7 – 14 day after noise. Here, we examined OHC ribbons immediately following prolonged noise exposure in apical regions of the cochlea. Although we found no significant differences, there was a slight downward trend in the number of CtBP2-expressing ribbons following noise in both genotypes. This slight decrease may suggest that modulation in the number of CtBP2 puncta may differ according to the duration of exposure or the post-exposure survival period. Loss of synaptic ribbons is also seen with aging and thought to be caused by cumulative damage, reduced afferent innervation, or changes in efferent input (Shi *et al*., 2015; Zhou *et al*., 2020; Liang *et al*., 2025).

Although the lack of OCM results in early-onset hearing loss and a greater sensitivity to noise damage, *Ocm^-/-^* mice have normal thresholds at least through the first 3 - 4 wks of age (Tong *et al*., 2016; Climer *et al*., 2021; Murtha *et al*., 2024). This period of normal thresholds in the absence of OCM suggests that there are alternative Ca^2+^ buffering mechanisms at play that regulate intracellular Ca^2+^ signals and maintain OHC function. The absence of OCM may alter one or more homeostatic control mechanisms involving Ca^2+^ influx and efflux. The primary sources of Ca^2+^ entry into OHCs are MET channels at the tips of stereocilia and voltage-gated Ca^2+^ channels (Ca_v_1.3) along the basolateral membrane (Knirsch *et al*., 2007; Pangrsic *et al*., 2018; Qiu & Muller, 2018). Several studies report that deletion of Ca_v_1.3 leads to an early loss of OHCs (Platzer *et al*.; Glueckert *et al*.; Engel *et al*.; Eckrich *et al*.). The primary source of Ca^2+^ extrusion in OHCs is the plasmalemmal Ca^2+^ pump isoform 2 (PMCA2), which is localized to OHC stereocilia (Dumont *et al*., 2001; Chen *et al*., 2012). Mutations in PMCA2 also lead to OHC loss and progressive hearing loss (Spiden *et al*., 2008; Bortolozzi *et al*., 2010). Disruption of any of the key proteins involved in OHC Ca^2+^ homeostasis leads to OHC dysfunction, loss, and subsequent hearing loss. We previously found that Ca_v_1.3 mRNA (*CACNA1D*), which serves as one of the main Ca^2+^ entry points in hair cells (Platzer *et al*., 2000; Michna *et al*., 2003), was down-regulated in *Ocm*^-/-^ mice (Yang *et al*., 2023). This downregulation might be a response to prevent Ca^2+^ overloading in OHCs and thus protect them from cytotoxicity. A similar compensatory mechanism may also exist in young adult OHCs to temporarily maintain normal cochlear function.

Since OHC Ca^2+^ buffering is dominated by OCM (Hackney *et al*., 2005), in the absence of OCM, other mobile buffers such as calbindin-D28k or αPV might substitute for its absence. However, several studies have shown changes in other mobile EF-hand CaBPs are not likely responsible for modulating intracellular Ca^2+^ levels (Pangrsic *et al*., 2015; Murtha *et al*., 2022; Yang *et al*., 2023). In cochlear hair cells, mitochondria can also modulate intracellular Ca^2+^ homeostasis via the mitochondrial Ca^2+^ uniporter (MCU) and by extruding Ca^2+^ back to the cytosol through the sodium-calcium exchanger (NCLX). Wang et al. (2018) has shown that MCU expression increased after noise exposure while the expression of NCLX decreased in a time- and exposure-dependent manner. Thus, OHC mitochondria could compensate for the absence of OCM and temporarily maintain the function of OHCs. However, long-term mitochondrial Ca^2+^ overload can lead to mitochondrial stress, induce OHC death, and subsequent hearing loss (Vicente-Torres & Schacht, 2006). This latter scenario might explain the early, progressive OHC loss observed in *Ocm*^-/-^ mice. In a recent report (Murtha *et al*., 2024), we suggested that the loss of OCM leads to mitochondrial degradation due to prolonged Ca^2+^ overload, which made OHCs more prone to degeneration after noise. Murtha et al (2024) observed that *Ocm*^-/-^ OHCs were more vulnerable to damage and loss as soon as 48 hrs after intense noise (106 dB SPL, 2 hrs). As in the present study, *Ocm*^-/-^ mice showed significantly greater threshold shifts and significant OHC loss, especially in basal regions of the cochlea compared to *Ocm*^+/+^ mice. Murtha et al (2024) reported that a higher level of OHC loss coincided with alterations in OHC mitochondrial morphology. Thus, OCM may protect cells against Ca^2+^-induced cell death by maintaining Ca^2+^ homeostasis. Both OCM and mitochondria are likely to determine the ability of OHCs to compensate for large increases in Ca^2+^ after acoustic overexposure. In line with other reports (Fettiplace & Nam, 2019), our results suggest that the vulnerability of OHCs depends upon their ability to maintain Ca^2+^ homeostasis and in this way, OCM is protective. Furthermore, the presence of threshold shifts despite the lack of OHC loss after moderate noise exposure, suggests OCM may play a role in delaying cellular senescence.

As in our previous studies in prehearing mice (Yang *et al*., 2023), we found that the expression of P2X2 purinergic receptors was upregulated in young adult *Ocm*^-/-^ mice compared to controls. The upregulation of purinergic receptors could be a compensatory mechanism for Ca^2+^ overload. In mature hair cells, P2X2 is a modulator of hair cell mechanotransduction, which plays an essential role in hair cell tolerance to noise (Mittal *et al*., 2016). In agreement with our study, others have reported that noise exposure leads to an upregulation of *P2RX2* mRNA and protein levels in the organ of Corti (Wang *et al*., 2003; Telang *et al*., 2010). The high expression of purinergic receptors, with an increase in local ATP release in the endolymph and perilymph (Munoz *et al*., 1995; Telang *et al*., 2010), may provide a shunt capable of regulating the electrical potential across the endolylmphatic surface of hair cells (Thorne *et al*., 2004). The mutation of *P2RX2* in both humans and mice gives rise to progressive hearing loss and increased sensitivity to noise-induced permanent threshold shifts (Yan *et al*., 2013). Furthermore, a recent study revealed an increase in P2Y purinergic response in supporting cells from the aged cochlea (Hool *et al*., 2023), indicating a possible protective mechanism that prevents further damage to the sensory epithelium. Since purinergic signaling also contributes to inflammation and immune responses (Burnstock, 2016; Liu *et al*., 2023), the long-term upregulation of purinergic receptors in OHCs from *Ocm*^-/-^ mice could ultimately damage hearing.

Our results further demonstrate the importance of OCM as a specialized Ca^2+^ buffer in OHCs. The homeostatic control of intracellular Ca^2+^ is likely to contribute to the sensitivity of the OHCs to cochlear injury (Mammano, 2011). Ca^2+^ signaling plays fundamental roles in cochlear development and function, including mechanotransduction, neurotransmitter release, innervation, and motility (Ceriani *et al*., 2025). We recently demonstrated that OHCs in the prehearing cochlea of *Ocm*^-/-^ mice have altered Ca^2+^ signaling and delayed maturation of innervation (Yang *et al*., 2023). Aberrant Ca^2+^ dynamics in hair cells has been shown to lead to cochlear pathology (Richard *et al*., 2023) and increased OHC loss in *Ocm*^-/-^ mice (Tong *et al*., 2016; Climer *et al*., 2021). Noise exposure, a common cause of hearing loss, leads to increases in the concentration of cytosolic free Ca^2+^ (Fridberger *et al*., 1998a; Zuo *et al*., 2008). Ca^2+^ overload in sensory hair cells can activate regulated cell-death pathways, which in turn, could exacerbate inflammatory responses (Fridberger *et al*., 1998a; Bienkowski *et al*., 2000; Orrenius *et al*., 2003; Esterberg *et al*., 2013; Fu *et al*., 2021). Thus, tightly regulating Ca^2+^ signaling through specialized mobile CaBPs such as OCM is essential for OHC function and survival and minimizes the consequences of noise-induced cochlear insults.

In summary, OCM protects OHCs from early cochlear pathology and from acoustic overstimulation. As schematized in **Figure 9**, wild-type OHCs repond to moderate levels of noise by inducing higher levels of purinergic receptor expression and higher levels of intracellular Ca^2+^, which lead to temporary changes in OHC amplification and ABR wave I amplitude and latency. The lack of OCM induces chronic dysregulation of Ca^2+^ represented by increased ATP-induced Ca^2+^ activity, higher expression of ATP receptors in OHCs, and abnormal ABR wave I responses. Adult *Ocm*^-/-^ mice show a noise-exposed phenotype, which likely leads to increased susceptibility to noise damage. We propose that as a CaBP, OCM is an important regulator of homeostatic Ca^2+^ signaling in OHCs and protects OHC from the deleterious consequences of Ca^2+^ overload.

**Graphical Abstract or Figure 9.**
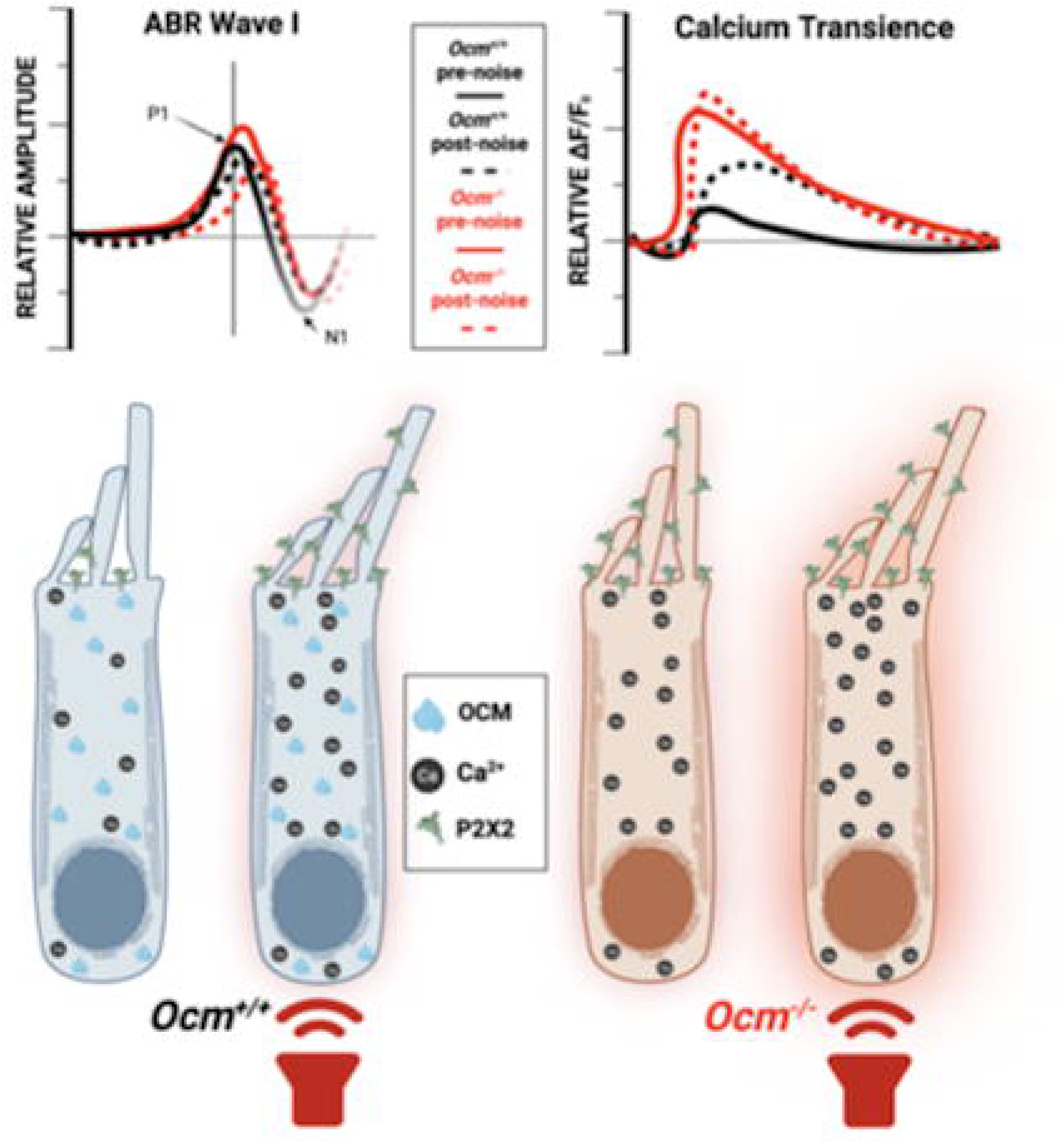
Schematic diagram representing altered ABR waveforms and calcium transience levels in *Ocm*^-/-^ OHCs and *Ocm*^+/+^ OHCs with moderate (95 dB SPL) noise exposure. In *Ocm*^+/+^ mice, moderate noise induces slight latency and amplitude shifts, increased purinergic receptor expression and increased calcium signaling. *Ocm*^-/-^ OHCs exhibit elevated calcium signaling pre-noise exposure and exhibit more dramatic latency and amplitude shifts post-noise exposure with no significant change in purinergic receptor expression after moderate noise exposure.

## Footnotes

1 https://www.masseyeandear.org/research/otolaryngology/eaton-peabody-laboratories/histology-core

## References

Bienkowski, P., Scinska, A., Kostowski, W., Koros, E. & Kukwa, A. (2000) Ototoxic mechanism of aminoglycoside antibiotics--role of glutaminergic NMDA receptors. Pol Merkur Lekarski, 9, 713–715.

Bortolozzi, M., Brini, M., Parkinson, N., Crispino, G., Scimemi, P., De Siati, R.D., Di Leva, F., Parker, A., Ortolano, S., Arslan, E., Brown, S.D., Carafoli, E. & Mammano, F. (2010) The novel PMCA2 pump mutation Tommy impairs cytosolic calcium clearance in hair cells and links to deafness in mice. J Biol Chem, 285, 37693–37703.

Buchwald, J. & Huang, C. (1975) Far-field acoustic response: origins in the cat. Science, 189, 382–384.

Burnstock, G. (2016) P2X ion channel receptors and inflammation. Purinergic Signal, 12, 59–67.

Ceriani, F. (2024) abrWaveAnalyser, Zenodo.

Ceriani, F., Hendry, A., Jeng, J.Y., Johnson, S.L., Stephani, F., Olt, J., Holley, M.C., Mammano, F., Engel, J. & Kros, C.J. (2019) Coordinated calcium signalling in cochlear sensory and non-sensory cells refines afferent innervation of outer hair cells. The EMBO journal, 38.

Ceriani, F., Kc, W., Sl, J., Cj, K. & W, M. (2025) Mechanisms driving the functional maturation of the developing mammalian auditory pathway. Current Topics in Developmental Biology.

Chen, F.Q., Zheng, H.W., Schacht, J. & Sha, S.H. (2013) Mitochondrial peroxiredoxin 3 regulates sensory cell survival in the cochlea. PLoS One, 8, e61999.

Chen, G., Racay, P., Bichet, S., Celio, M.R., Eggli, P. & Schwaller, B. (2006) Deficiency in parvalbumin, but not in calbindin D-28k upregulates mitochondrial volume and decreases smooth endoplasmic reticulum surface selectively in a peripheral, subplasmalemmal region in the soma of Purkinje cells. Neuroscience, 142, 97–105.

Chen, J., Chu, H., Xiong, H., Chen, Q., Zhou, L., Bing, D., Liu, Y., Gao, Y., Wang, S., Huang, X. & Cui, Y. (2012) Expression patterns of Ca(V)1.3 channels in the rat cochlea. Acta Biochim Biophys Sin (Shanghai), 44, 513–518.

Climer, L.K., Cox, A.M., Reynolds, T.J. & Simmons, D.D. (2019) Oncomodulin: The Enigmatic Parvalbumin Protein. Front Mol Neurosci, 12, 235.

Climer, L.K., Hornak, A.J., Murtha, K., Yang, Y., Cox, A.M., Simpson, P.L., Le, A. & Simmons, D.D. (2021) Deletion of Oncomodulin Gives Rise to Early Progressive Cochlear Dysfunction in C57 and CBA Mice. Frontiers in Aging Neuroscience, 13.

Cox, B.C., Liu, Z., Lagarde, M.M.M. & Zuo, J. (2012) Conditional Gene Expression in the Mouse Inner Ear Using Cre-loxP. Journal of the Association for Research in Otolaryngology, 13, 295–322.

Dallos, P. (1992) The active cochlea. J Neurosci, 12, 4575–4585.

Dumont, R.A., Lins, U., Filoteo, A.G., Penniston, J.T., Kachar, B. & Gillespie, P.G. (2001) Plasma membrane Ca2+-ATPase isoform 2a is the PMCA of hair bundles. J Neurosci, 21, 5066–5078.

Eckrich, S., Hecker, D., Sorg, K., Blum, K., Fischer, K., Munkner, S., Wenzel, G., Schick, B. & Engel, J. (2019) Cochlea-Specific Deletion of Cav1.3 Calcium Channels Arrests Inner Hair Cell Differentiation and Unravels Pitfalls of Conditional Mouse Models. Front Cell Neurosci, 13, 225.

Engel, J., Braig, C., Rüttiger, L., Kuhn, S., Zimmermann, U., Blin, N., Sausbier, M., Kalbacher, H., Münkner, S., Rohbock, K., Ruth, P., Winter, H. & Knipper, M. (2006) Two classes of outer hair cells along the tonotopic axis of the cochlea. Neuroscience, 143, 837–849.

Esterberg, R., Hailey, D.W., Coffin, A.B., Raible, D.W. & Rubel, E.W. (2013) Disruption of intracellular calcium regulation is integral to aminoglycoside-induced hair cell death. J Neurosci, 33, 7513–7525.

Fernandez, E.A., Ohlemiller, K.K., Gagnon, P.M. & Clark, W.W. (2010) Protection against noise-induced hearing loss in young CBA/J mice by low-dose kanamycin. J Assoc Res Otolaryngol, 11, 235–244.

Fettiplace, R. & Nam, J.-H. (2019) Tonotopy in calcium homeostasis and vulnerability of cochlear hair cells. Hearing research, 376, 11–21.

Fridberger, Flock, Å., Ulfendahl, M. & Flock, B. (1998a) Acoustic overstimulation increases outer hair cell Ca2+ concentrations and causes dynamic contractions of the hearing organ. Proceedings of the National Academy of Sciences, 95, 7127–7132.

Fridberger, A., Flock, A., Ulfendahl, M. & Flock, B. (1998b) Acoustic overstimulation increases outer hair cell Ca2+ concentrations and causes dynamic contractions of the hearing organ. Proceedings of the National Academy of Sciences of the United States of America, 95, 7127–7132.

Fridberger, A. & Ulfendahl, M. (1996) Acute mechanical overstimulation of isolated outer hair cells causes changes in intracellular calcium levels without shape changes. Acta Otolaryngol, 116, 17–24.

Fu, X., Wan, P., Li, P., Wang, J., Guo, S., Zhang, Y., An, Y., Ye, C., Liu, Z., Gao, J., Yang, J., Fan, J. & Chai, R. (2021) Mechanism and Prevention of Ototoxicity Induced by Aminoglycosides. Front Cell Neurosci, 15, 692762.

Glueckert, R., Wietzorrek, G., Kammen-Jolly, K., Scholtz, A., Stephan, K., Striessnig, J. & Schrott-Fischer, A. (2003) Role of class D L-type Ca2+ channels for cochlear morphology. Hear Res, 178, 95–105.

Hackney, C.M., Mahendrasingam, S., Penn, A. & Fettiplace, R. (2005) The concentrations of calcium buffering proteins in mammalian cochlear hair cells. J Neurosci, 25, 7867–7875.

Hirose, K. & Liberman, M.C. (2003) Lateral wall histopathology and endocochlear potential in the noise-damaged mouse cochlea. J Assoc Res Otolaryngol, 4, 339–352.

Hool, S.A., Jeng, J.Y., Jagger, D.J., Marcotti, W. & Ceriani, F. (2023) Age-related changes in P2Y receptor signalling in mouse cochlear supporting cells. J Physiol, 601, 4375–4395.

Huang, L.-C., Barclay, M., Lee, K., Peter, S., Housley, G.D., Thorne, P.R. & Montgomery, J.M. (2012) Synaptic profiles during neurite extension, refinement and retraction in the developing cochlea. Neural Development, 7, 1–17.

Jeng, J.-Y., Carlton, A.J., Johnson, S.L., Brown, S.D.M., Holley, M.C., Bowl, M.R. & Marcotti, W. (2021) Biophysical and morphological changes in inner hair cells and their efferent innervation in the ageing mouse cochlea. The Journal of Physiology, 599, 269–287.

Jeng, J.Y., Ceriani, F., Hendry, A., Johnson, S.L., Yen, P., Simmons, D.D., Kros, C.J. & Marcotti, W. (2020) Hair cell maturation is differentially regulated along the tonotopic axis of the mammalian cochlea. J Physiol, 598, 151–170.

Johnson, S.L., Safieddine, S., Mustapha, M. & Marcotti, W. (2019) Hair Cell Afferent Synapses: Function and Dysfunction. Cold Spring Harb Perspect Med, 9.

Kidd Iii, A.R. & Bao, J. (2012) Recent advances in the study of age-related hearing loss: a mini-review. Gerontology, 58, 490–496.

Knirsch, M., Brandt, N., Braig, C., Kuhn, S., Hirt, B., Munkner, S., Knipper, M. & Engel, J. (2007) Persistence of Ca(v)1.3 Ca2+ channels in mature outer hair cells supports outer hair cell afferent signaling. J Neurosci, 27, 6442–6451.

Kujawa, S.G. & Liberman, M.C. (2009) Adding insult to injury: cochlear nerve degeneration after “temporary” noise-induced hearing loss. J Neurosci, 29, 14077–14085.

Lavinsky, J., Crow, A.L., Pan, C., Wang, J., Aaron, K.A., Ho, M.K., Li, Q., Salehide, P., Myint, A., Monges-Hernadez, M., Eskin, E., Allayee, H., Lusis, A.J. & Friedman, R.A. (2015) Genome-wide association study identifies nox3 as a critical gene for susceptibility to noise-induced hearing loss. PLoS Genet, 11, e1005094.

Liang, C., Zhai, T.Y., Chen, J., Fang, S., Zhu, Y., Liu, L.M., Yu, N. & Zhao, H.B. (2025) ATP-gated P2×7 receptors express at type II auditory nerves and required for efferent hearing control and noise protection. Proc Natl Acad Sci U S A, 122, e2421995122.

Liberman, M. & Mulroy, M. (1982) Acute and chronic effects of acoustic trauma: cochlear pathology and auditory-nerve pathophysiology. In Rp, H. D. H. R. S. (eds) New Perspectives on Noise-Induced Hearing Loss. Raven Press, New York, pp. 105–135.

Liberman, M.C. & Kujawa, S.G. (2017) Cochlear synaptopathy in acquired sensorineural hearing loss: Manifestations and mechanisms. Hear Res, 349, 138–147.

Liu, J.P., Liu, S.C., Hu, S.Q., Lu, J.F., Wu, C.L., Hu, D.X. & Zhang, W.J. (2023) ATP ion channel P2X purinergic receptors in inflammation response. Biomed Pharmacother, 158, 114205.

Liu, X., Bulgakov, O.V., Darrow, K.N., Pawlyk, B., Adamian, M., Liberman, M.C. & Li, T. (2007) Usherin is required for maintenance of retinal photoreceptors and normal development of cochlear hair cells. Proc Natl Acad Sci U S A, 104, 4413–4418.

Liu, Y. & Fechter, L.D. (1997) Toluene disrupts outer hair cell morphometry and intracellular calcium homeostasis in cochlear cells of guinea pigs. Toxicol Appl Pharmacol, 142, 270–277.

Maison, S.F., Liu, X.P., Eatock, R.A., Sibley, D.R., Grandy, D.K. & Liberman, M.C. (2012) Dopaminergic signaling in the cochlea: receptor expression patterns and deletion phenotypes. J Neurosci, 32, 344–355.

Mammano, F. (2011) Ca2+ homeostasis defects and hereditary hearing loss. BioFactors, 37, 182–188.

Marcotti, W. & Kros, C.J. (1999) Developmental expression of the potassium current IK,n contributes to maturation of mouse outer hair cells. The Journal of physiology, 520 **Pt** 3, 653–660.

Mehraei, G., Hickox, A.E., Bharadwaj, H.M., Goldberg, H., Verhulst, S., Liberman, M.C. & Shinn-Cunningham, B.G. (2016) Auditory Brainstem Response Latency in Noise as a Marker of Cochlear Synaptopathy. J Neurosci, 36, 3755–3764.

Melgar-Rojas, P., Alvarado, J.C., Fuentes-Santamaria, V., Gabaldon-Ull, M.C. & Juiz, J.M. (2015) Validation of Reference Genes for RT-qPCR Analysis in Noise-Induced Hearing Loss: A Study in Wistar Rat. PLoS One, 10, e0138027.

Michna, M., Knirsch, M., Hoda, J.C., Muenkner, S., Langer, P., Platzer, J., Striessnig, J. & Engel, J. (2003) Cav1.3 (alpha1D) Ca2+ currents in neonatal outer hair cells of mice. J Physiol, 553, 747–758.

Mittal, R., Grati, M., Sedlacek, M., Yuan, F., Chang, Q., Yan, D., Lin, X., Kachar, B., Farooq, A., Chapagain, P., Zhang, Y. & Liu, X.Z. (2016) Characterization of ATPase Activity of P2RX2 Cation Channel. Front Physiol, 7, 186.

Mulvaney, J. & Dabdoub, A. (2012) Atoh1, an Essential Transcription Factor in Neurogenesis and Intestinal and Inner Ear Development: Function, Regulation, and Context Dependency. Journal of the Association for Research in Otolaryngology, 13, 281–293.

Munoz, D.J., Thorne, P.R., Housley, G.D. & Billett, T.E. (1995) Adenosine 5’-triphosphate (ATP) concentrations in the endolymph and perilymph of the guinea-pig cochlea. Hear Res, 90, 119–125.

Murtha, K.E., Sese, W.D., Sleiman, K., Halpage, J., Padyala, P., Yang, Y., Hornak, A.J. & Simmons, D.D. (2024) Absence of oncomodulin increases susceptibility to noise-induced outer hair cell death and alters mitochondrial morphology. Front Neurol, 15, 1435749.

Murtha, K.E., Yang, Y., Ceriani, F., Jeng, J.Y., Climer, L.K., Jones, F., Charles, J., Devana, S.K., Hornak, A.J., Marcotti, W. & Simmons, D.D. (2022) Oncomodulin (OCM) uniquely regulates calcium signaling in neonatal cochlear outer hair cells. Cell Calcium, 105, 102613.

Orrenius, S., Zhivotovsky, B. & Nicotera, P. (2003) Regulation of cell death: the calcium-apoptosis link. Nat Rev Mol Cell Biol, 4, 552–565.

Pangrsic, T., Gabrielaitis, M., Michanski, S., Schwaller, B., Wolf, F., Strenzke, N. & Moser, T. (2015) EF-hand protein Ca2+ buffers regulate Ca2+ influx and exocytosis in sensory hair cells. Proc Natl Acad Sci U S A, 112, E1028–1037.

Pangrsic, T., Singer, J.H. & Koschak, A. (2018) Voltage-Gated Calcium Channels: Key Players in Sensory Coding in the Retina and the Inner Ear. Physiol Rev, 98, 2063–2096.

Platzer, J., Engel, J., Schrott-Fischer, A., Stephan, K., Bova, S., Chen, H., Zheng, H. & Striessnig, J. (2000) Congenital deafness and sinoatrial node dysfunction in mice lacking class D L-type Ca2+ channels. Cell, 102, 89–97.

Qiu, X. & Muller, U. (2018) Mechanically Gated Ion Channels in Mammalian Hair Cells. Front Cell Neurosci, 12, 100.

Richard, E.M., Maurice, T. & Delprat, B. (2023) Calcium signaling and genetic rare diseases: An auditory perspective. Cell Calcium, 110, 102702.

Schmittgen, T.D. & Livak, K.J. (2008) Analyzing real-time PCR data by the comparative C(T) method. Nat Protoc, 3, 1101–1108.

Shi, L., Liu, K., Wang, H., Zhang, Y., Hong, Z., Wang, M., Wang, X., Jiang, X. & Yang, S. (2015) Noise induced reversible changes of cochlear ribbon synapses contribute to temporary hearing loss in mice. Acta Oto-Laryngologica, 135, 1093–1102.

Simmons, D.D., Tong, B., Schrader, A.D. & Hornak, A.J. (2010) Oncomodulin identifies different hair cell types in the mammalian inner ear. J Comp Neurol, 518, 3785–3802.

Spiden, S.L., Bortolozzi, M., Di Leva, F., de Angelis, M.H., Fuchs, H., Lim, D., Ortolano, S., Ingham, N.J., Brini, M., Carafoli, E., Mammano, F. & Steel, K.P. (2008) The novel mouse mutation Oblivion inactivates the PMCA2 pump and causes progressive hearing loss. PLoS Genet, 4, e1000238.

Stamper, G.C. & Johnson, T.A. (2015) Auditory function in normal-hearing, noise-exposed human ears. Ear Hear, 36, 172–184.

Telang, R.S., Paramananthasivam, V., Vlajkovic, S.M., Munoz, D.J., Housley, G.D. & Thorne, P.R. (2010) Reduced P2x(2) receptor-mediated regulation of endocochlear potential in the ageing mouse cochlea. Purinergic Signal, 6, 263–272.

Thorne, P.R., Munoz, D.J. & Housley, G.D. (2004) Purinergic modulation of cochlear partition resistance and its effect on the endocochlear potential in the Guinea pig. J Assoc Res Otolaryngol, 5, 58–65.

Tong, B., Hornak, A.J., Maison, S.F., Ohlemiller, K.K., Liberman, M.C. & Simmons, D.D. (2016) Oncomodulin, an EF-Hand Ca 2+ Buffer, Is Critical for Maintaining Cochlear Function in Mice. The Journal of Neuroscience, 36, 1631–1635.

Vicente-Torres, M.A. & Schacht, J. (2006) A BAD link to mitochondrial cell death in the cochlea of mice with noise-induced hearing loss. J Neurosci Res, 83, 1564–1572.

Wang, J.C., Raybould, N.P., Luo, L., Ryan, A.F., Cannell, M.B., Thorne, P.R. & Housley, G.D. (2003) Noise induces up-regulation of P2X2 receptor subunit of ATP-gated ion channels in the rat cochlea. Neuroreport, 14, 817–823.

Wang, X., Zhu, Y., Long, H., Pan, S., Xiong, H., Fang, Q., Hill, K., Lai, R., Yuan, H. & Sha, S.H. (2018) Mitochondrial Calcium Transporters Mediate Sensitivity to Noise-Induced Losses of Hair Cells and Cochlear Synapses. Front Mol Neurosci, 11, 469.

Wood, M.B., Nowak, N., Mull, K., Goldring, A., Lehar, M. & Fuchs, P.A. (2021) Acoustic Trauma Increases Ribbon Number and Size in Outer Hair Cells of the Mouse Cochlea. JARO, 22, 19–31.

Yan, D., Zhu, Y., Walsh, T., Xie, D., Yuan, H., Sirmaci, A., Fujikawa, T., Wong, A.C., Loh, T.L., Du, L., Grati, M., Vlajkovic, S.M., Blanton, S., Ryan, A.F., Chen, Z.Y., Thorne, P.R., Kachar, B., Tekin, M., Zhao, H.B., Housley, G.D., King, M.C. & Liu, X.Z. (2013) Mutation of the ATP-gated P2X(2) receptor leads to progressive hearing loss and increased susceptibility to noise. Proc Natl Acad Sci U S A, 110, 2228–2233.

Yang, H., Xie, X., Deng, M., Chen, X. & Gan, L. (2010) Generation and characterization of Atoh1-Cre knock-in mouse line. Genesis, 48, 407–413.

Yang, Y., Murtha, K., Climer, L.K., Ceriani, F., Thompson, P., Hornak, A.J., Marcotti, W. & Simmons, D.D. (2023) Oncomodulin regulates spontaneous calcium signalling and maturation of afferent innervation in cochlear outer hair cells. The Journal of Physiology.

Zhou, H., Qian, X., Xu, N., Zhang, S., Zhu, G., Zhang, Y., Liu, D., Cheng, C., Zhu, X., Liu, Y., Lu, L., Tang, J., Chai, R. & Gao, X. (2020) Disruption of Atg7-dependent autophagy causes electromotility disturbances, outer hair cell loss, and deafness in mice. Cell Death Dis, 11, 913.

Zuo, H., Cui, B., She, X. & Wu, M. (2008) Changes in Guinea pig cochlear hair cells after sound conditioning and noise exposure. J Occup Health, 50, 373–379.

